# Genetic determinants of plasma protein levels in the Estonian population

**DOI:** 10.1101/2023.05.30.542983

**Authors:** Anette Kalnapenkis, Maarja Jõeloo, Kaido Lepik, Viktorija Kukuškina, Mart Kals, Kaur Alasoo, Estonian Biobank Research Team, Reedik Mägi, Tõnu Esko, Urmo Võsa

## Abstract

The proteome holds great potential as an intermediate layer between the genome and phenome. Previous protein quantitative trait locus studies have focused mainly on describing the effects of common genetic variations on the proteome. Here, we assessed the impact of the common and rare genetic variations as well as the copy number variants (CNVs) on 326 plasma proteins measured in up to 500 individuals. We identified 184 *cis* and 94 *trans* signals for 157 protein traits, which were further fine-mapped to credible sets for 101 *cis* and 87 *trans* signals for 151 proteins. Rare genetic variation contributed to the levels of 7 proteins, with 5 *cis* and 14 *trans* associations. CNVs were associated with the levels of 11 proteins (7 *cis* and 5 *trans*), examples including a 3q12.1 deletion acting as a hub for multiple *trans* associations; and a CNV overlapping *NAIP*, a sensor component of the NAIP-NLRC4 inflammasome which is affecting pro-inflammatory cytokine interleukin 18 levels. In summary, this work presents a comprehensive resource of genetic variation affecting the plasma protein levels and provides the interpretation of identified effects.

## Introduction

During the last decade, genome-wide association studies (GWASs) have successfully linked genetic variants to complex traits [1]. However, the mechanisms underlying many of these associations often remain unknown, as most of the associated genetic variants are located in non-coding regions of the genome, suggesting that they have regulatory effects on phenotypes [2]. To fill this knowledge gap, molecular traits are routinely used as intermediate phenotypes in association studies. The study of molecular phenotypes enables the assessment of the direct effects of genetic variants on, for example, the alteration of protein levels, and the potential underlying molecular mechanisms and links to endpoint phenotypes.

Proteins are functional products of the genome that provide insight about the normal processes of organisms; in addition, alterations in their levels are indicators of changes in disease status [3]. Recent technological advancements, including the development of multiplex immunoassays and aptamer assays, have provided opportunities for the measurement of thousands of plasma- and serum-based protein levels [4–8].

The genetic backgrounds of protein levels are uncovered through the linking of these levels to genetic variability via protein quantitative trait locus (pQTL) analysis. Many recent pQTL studies have been large-scale [4–8], with the largest of them including 54,306 individuals from the UK Biobank [9]. Their primary focus has been the identification of common [minor allele frequency (MAF) > 0.01] variants affecting inter-individual protein variability, but Sun et al. [9] reported that approximately 5.6% (570/10,248) and 1.5% (155/10,248) of the variants with primary associations had MAFs < 0.01 and < 0.005, respectively. In addition, the focus has been shifting toward the identification of associations with rare (MAF < 0.01) variants, using gene-based methods [10–14]. For example, a recent landmark study conducted on the Icelandic population revealed 18,084 genetic associations with protein levels, 19% of which were with rare variants [8]. Investigation of the effects of other structural variants, such as copy number variants (CNVs), on protein levels has thus far been limited [15].

The combined examination of pQTL and GWAS results for disease phenotypes can lead to the validation and prioritisation of new and existing drug targets, and the identification of clinically relevant biomarkers. Ferkingstad et al. [8] found that 12% of 45,334 lead associations in the GWAS Catalog were with variants in high linkage disequilibrium (LD) with pQTLs. The application of Mendelian randomisation (MR) and colocalisation analysis to biomedical data for the identification of links between pQTLs and diseases enables the evaluation of the causality between protein levels and disease risk and the identification of potential disease pathways, respectively. Zheng et al. [16] used MR and colocalisation analysis to examine associations of 1,002 plasma proteins with 153 diseases and 72 disease-related risk factors, and identified 413 protein–trait associations supported by MR, 130 (31.5%) of which were not supported by the colocalisation analysis. This example highlights the importance of intersecting the results from both analyses [17].

Here, we integrated dense whole-genome sequencing (WGS) data to study the genetic contributions of rare and common variants to 326 plasma protein levels in the Estonian Biobank cohort (Fig 1). We examined the effects of single nucleotide polymorphisms (SNPs) and common CNVs on the inter-individual protein variability, and identified several proteins that were affected by the latter. To assess the overlap of local (*cis*) and distal (*trans*) pQTL effects with gene expression levels, we conducted comprehensive colocalisation analyses with expression quantitative trait loci (eQTLs) and splicing QTLs using data from various tissues from the eQTL Catalogue [18].

**Figure 1.**
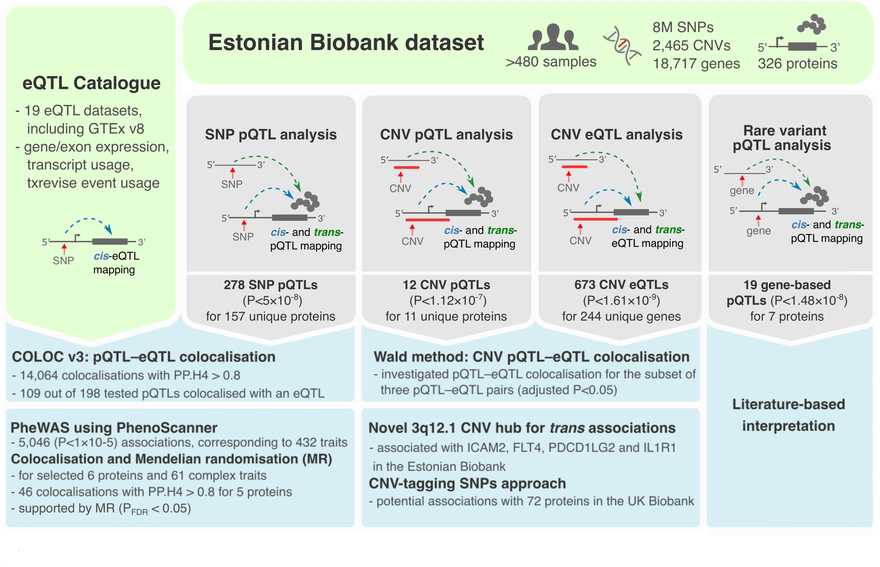
Overview of the main analyses conducted in this study.

## Material and methods

### Study samples

The Estonian Biobank (EstBB) cohort consists of more than 200,000 Estonian volunteers aged ≥ 18 years, representing about 20% of the Estonian adult population, detailed information on the enrollment process and data collection is described in the Leitsalu et al. study [19]. Genotype data are available for all gene donors in this cohort. For a subcohort of 500 individuals [52.8% females and 47.2% males, mean age 54 (standard deviation 14.0) years], WGS, RNA sequencing and Olink proteomics data from the same timepoint are available. The WGS dataset was generated in 2015. Sample collection for RNA sequencing and Olink proteomics was conducted in years 2011-2012. RNA sequencing was performed in years 2015-2016 and protein levels were measured in year 2017. The activities of the EstBB are regulated by the Human Genes Research Act, which was adopted in year 2000 specifically for the operations of the EstBB. All participants have signed a broad consent form to allow researchers to use their genomics and health data for studies upon approval by the Estonian Committee on Bioethics and Human Research. Individual level data analysis for this project was carried out under approval 1.1-12/624 from the Estonian Committee on Bioethics and Human Research (Estonian Ministry of Social Affairs) and data extraction no. K29 from the Estonian Biobank. The current study was conducted using pseudonymised data.

### WGS data processing, variant calling and quality control

The 2,284 EstBB WGS samples were sequenced at the Genomics Platform of the Broad Institute (Cambridge, MA, USA). Sequenced data were jointly variant called and quality controlled as described by Mitt et al. [20]; and the final WGS sample set was derived from 2,244 individuals. We excluded multiallelic sites and genetic variants, based on quality/depth < 2, Hardy–Weinberg equilibrium test failure (*P* > 1×10^−9^), and call rate < 90%. Data from individuals with available proteomics data (*n* = 500) were retained for further analyses.

### CNV detection and quality control

The Genome STRiP pipeline (version 2.00.1611) [21] was applied to detect CNVs from aligned sequencing reads (in BAM format) for 2,284 samples as described by Lepamets et al. [22]. In brief, CNV sites were identified and genotyped in five batches. After the exclusion of samples with excessive numbers of calls, the batches were combined and duplicate calls were merged. Low-quality calls and sites with call rates <90% were excluded. We restricted the final dataset to deletions longer than 1,000 bp and duplications longer than 2,000 bp. The final sample set contained 51,026 CNV sites from 2,230 individuals. Data from individuals with available proteomics data (*n* = 500) were retained for further analyses.

### Measurement of plasma protein levels

Plasma concentrations in EDTA plasma samples from 500 Estonian Biobank donors were measured using four arrays with 92 protein targets each [ProSeek Cardiovascular Disease (CVD) II and III, Inflammation and Oncology II; Olink Biosciences, Uppsala, Sweden; S1 Table]. The procedure is described in detail elsewhere [23], and a technical white paper with additional information is available at the manufacturer’s website (https://www.olink.com). The native Olink data consisted of qPCR cycle threshold values corrected for extension control, followed by inter-plate control and the application of a correction factor predetermined by a negative control signal. The measurements were given at a natural logarithmic scale as normalised protein expression levels, a relative quantification unit. As part of the quality control, we excluded individual samples that did not pass the Olink internal quality control system. Final sample sizes per array ranged from 488 to 497, and the samples were measured in six batches. For arrays in which <20% of samples had values below the limit of detection (LOD), protein level correction was performed by dividing the Olink-assigned LOD value by 2, as done in the SCALLOP CVD-I project [6]. A total of 341 protein traits (326 unique proteins, as 14 proteins were measured by more than one array) passed quality control and were retained for further analyses (S1 Table).

### RNA sequencing data

RNA was extracted from samples in thawed Tempus tubes using TRIzol reagent (Invitrogen, Waltham, MA, USA) and further purified using an RNeasy Mini Kit (Qiagen, Hilden, Germany). Globin mRNA was depleted using GLOBINclear Kit (Invitrogen, Waltham, MA, USA). RNA quality was checked using electrophoresis (Agilent 2200 TapeStation; Agilent Technologies, Santa Clara, CA, USA). Sequencing libraries were prepared using 200 ng RNA according to the Illumina TruSeq stranded mRNA protocol. RNA sequencing was performed at the Estonian Genome Centre Core Facility using paired-end 50-bp sequencing technology (Illumina, San Diego, CA, USA), according to the manufacturer’s specifications.

Adapters and leading and trailing bases with a quality score were removed using Trimmomatic (version 0.36) [24]. Quality control was done with FastQC (version 0.11.2) [25]. Reads were mapped to human genome reference version GRCh37.p13 with STAR (version 2.4.2a) [26]. Reads that mapped to each genomic feature were counted with STAR using the same algorithm as default htseq-count. Raw RNA sequencing counts were normalised with the weighted trimmed mean of M-values [27] method from the edgeR R package (version 3.12.1) [28]. Detailed information regarding RNA sequencing data pre-processing is described in Lepik et al. [29]. The final gene expression measure was in logarithmed count per million. In total, 486 RNA sequencing samples overlapped with available proteomics data and were used for eQTL mapping.

### Genome-wide SNP pQTL discovery

Protein trait levels were rank-based inverse normal transformed. We regressed out the effects of age, sex, the season of sample collection, smoking status, blood sample processing time (days), plasma sample storage time (in days) and protein analysis batch using a custom R script. The residuals were used in a single-variant pQTL analysis performed with the EMMAX linear mixed model [30] and the EPACTS software (version 3.3.0, *q.emmax* function; https://genome.sph.umich.edu/wiki/EPACTS). To account for population structure, a kinship matrix was generated in EPACTS using genetic variants with MAF > 0.01 and call rate > 95%. Depending on the panel, we tested between 8,856,032 and 8,891,303 autosomal genetic variants against 341 plasma protein traits.

We classified associated variants into two categories based on their positions in relation to the protein-coding genes. We defined *cis*-pQTLs as SNPs located within 1Mb upstream or downstream of the transcription start sites (TSSs) of the corresponding protein-coding genes, and *trans*-pQTLs as SNPs located >1 Mb upstream or downstream of the TSS or on a different chromosome. Heterodimers were classified based on the protein subunit gene closest to the associated variant. In the case of proteins that were present on multiple panels, weaker signals were omitted from the analyses.

To retain independent signals, associated variants were clumped in PLINK (version 1.9) [31], using a 1 Mb window with the LD thresholds of *R^2^* = 0.1 and *P* < 5 × 10^−8^. To flag potential ‘pseudo-pQTL’ signals caused by the epitope effect, i.e. altered assay binding affinity due to a change in protein structure instead of an actual change in protein expression level, we followed the strategy described by Folkersen et al, 2020 [6]. Briefly, we determined whether any lead *cis* variant was a protein-altering variant (PAV) or in high LD (*R²* ≥ 0.8) with one, by using 2,230 WGS samples as the reference for the LD calculations (S2 Table). Missense, frameshift, splice donor region and stop gain variants were flagged as PAVs. Lead pQTL variants were queried for evidence of location in a regulatory region using RegulomeDB [32].

### Corresponding eQTL discovery

In order to overlap the genome-wide significant (*P* < 5 × 10^−8^) pQTLs with eQTLs, we used the RNA sequencing data from the overlapping samples of the same cohort [29]. We tested the eQTL effects on the genes encoding corresponding proteins by using a linear mixed model from EPACTS software (version 3.2.2) [30] with the same settings as for pQTL analysis. We included age, sex, body mass index, blood components (neutrophils, eosinophils, basophils, lymphocytes, monocytes, erythrocytes and thrombocytes) and RNA sequencing batch as covariates. To account for hidden batch effects on the gene expression, the first two principal components of the gene expression data were also included as covariates, as described in detail in Lepik et al. [29]. To correct for multiple testing, we adjusted P-values using false discovery rate (FDR) correction; eQTLs were considered as replicated at Benjamini-Hochberg FDR ≤ 0.05 and with concordant allelic direction with the pQTLs.

### Multiple testing correction for the pQTL analysis

From primary analyses, effects reaching per-protein genome-wide significance (*P* < 5 × 10^−8^) were interpreted. To also provide the more conservative results accounting for the number of tested proteins, we used a strategy described by Gao et al. and Kettunen et al. [33,34], which accommodates the correlation between protein levels. Four matrices corresponding to inverse normal transformed and covariate-adjusted protein levels from the Olink panels were merged. Only samples that passed quality control on every panel (*n* = 478) were included. The resulting matrix of standardised residuals was used in a principal components analysis implemented with the FactoMiner (version 1.41) [35] R package. As 181 principal components cumulatively explained >95% of the total variance in the proteomics data, the stricter significance threshold was set to 2.76 × 10^−10^ (5 × 10^−8^ / 181).

### Gene-based analysis of rare SNPs

Variants were annotated using the EPACTS ‘anno’ module (version 3.3.0; https://genome.sph.umich.edu/wiki/EPACTS) and GENCODE (version 14) [36] to ascertain their effects on protein sequences. A gene-based group file was generated with the inclusion of all nonsynonymous (missense and nonsense) variants in assigned genes. Only genes with more than two nonsynonymous variants were retained. We performed the gene-based SKAT test using the EMMAX *mmskat* function with adjustment for small sample size in EPACTS, using all variants with 0.000001% < MAF < 1%. Covariates included in the rare variant pQTL analysis were the same as described in the Methods section for Genome-wide SNP pQTL discovery. The results were corrected for multiple testing based on Bonferroni-corrected threshold of *P* < 1.48 × 10^−8^ [0.05 / (18,717 genes × 181 protein traits)]. Associations between genes and levels of proteins encoded on the same gene were classified as *cis*, and all other associations were classified as *trans*. Using the GeneMANIA database [37,38], we investigated whether the associated genes also had gene–gene functional interactions with corresponding protein-coding genes. For overlapping the rare variant pQTL associations with eQTL data, we performed an eQTL mapping with EPACTS software (version 3.2.2) using the same gene-based SKAT test as in rare variant pQTL mapping. Covariates included in the rare variant eQTL analysis were the same as described in the Methods section for Corresponding eQTL discovery. Similar to single variant eQTL analysis, to account for multiple testing, we adjusted *P*-values using false discovery rate (FDR) correction; rare variant eQTLs were considered as replicated at Benjamini-Hochberg FDR ≤ 0.05 and directionally concordant with the rare variant pQTLs.

### Fine-mapping analysis

We conducted a fine-mapping analysis to pinpoint causal variants for protein level–significant (*P* < 5 × 10^−8^) SNV-pQTL associations. We excluded the LTA and MICA-MICB proteins associated with variants in the major histocompatibility complex region on chromosome 6, due to the complexity of the associated *HLA* region. The fine-mapping procedure was based on the SuSiE ‘sum of single effects’ model [39,40] and was implemented using the *susie_suff_stat* function from susieR package (version 0.11.42). Fine-mapping pipeline was implemented in Nextflow [41] and some scripts were modified from the FINNGEN fine-mapping pipeline (https://github.com/FINNGEN/finemapping-pipeline). The SuSiE output contains single effect components, i.e., credible sets (CSs), with a >95% probability of including a variant with a non-zero causal effect. We used a default setting of 10 for the maximum number of causal variants regulating a protein, because Wang et al. has demonstrated it to be the optimal choice for the number of causal variants [39]. LDstore (version 2) [42] was used to generate an LD matrix for each locus.

### Replication of pQTLs

All significant lead variants from the pQTL discovery analyses were queried for previously published associations with protein levels in the PhenoScanner database (version 2) [43,44] using the Python application (https://github.com/phenoscanner/phenoscannerpy, query date 4 October 2021). This database contains results from large pQTL studies [4,45,46]. For variant matching between datasets, we created variant names that were concatenations of the corresponding chromosome, chromosome position (hg19), and alphabetically ordered alleles. To match UniProt IDs from the discovery analyses to PhenoScanner trait names, the IDs were converted to recommended HUGO Gene Nomenclature Committee gene names using the UniProt conversion tool (https://www.uniprot.org/uploadlists/, latest query date 11 October 2021). We performed additional replication analysis using Pietzner et al. dataset by querying their publicly available results with *P* < 0.05 [7]. The largest pQTL meta-analysis published to date (*n* = 30,931) [6] was conducted through the SCALLOP consortium and was not usable due to sample overlap with the current study. In order to ensure that each protein was represented by a single association, we restricted our comparisons to instances where either one subunit or the entire heterodimer complex was available. For instances where one protein was available multiple times, we conducted comparison with the association with the smallest *P*-value. To account for multiple testing, we adjusted *P*-values using false discovery rate (FDR) correction; pQTLs were considered as replicated at Benjamini-Hochberg FDR ≤ 0.05 and concordant allelic direction with the discovery pQTLs.

### Identification of relevant disease traits and molecular QTLs

To identify complex traits and diseases associated with the top pQTLs, we conducted a phenome-wide association analysis (PheWAS) by querying the lead variants from primary pQTL mapping and their proxies against the PhenoScanner database (version 2) [43,44]. Duplicate associations happening due to data resource overlap were removed. We considered only PhenoScanner associations with *P* < 1 × 10^−5^. Specifically, we sought to identify pQTLs associated with disease traits, methylation quantitative trait loci (meQTLs), histone modifications and metabolite quantitative trait loci (mQTLs), as well as percent-spliced-in (PSI) associations. We also searched for significant protein genes on a druggable genome list [47] and the drugs that interact with them [48]. For a subset of pQTLs we selected for in-depth analyses by coloc and Mendelian randomisation, an additional PheWAS was conducted with the Medical Research Council (MRC) Integrative Epidemiology Unit (IEU) OpenGWAS database [49]. This was done to extract region-wide associations, irrespective of association *P*-value.

### Colocalisation analysis

The colocalisation analyses between pQTLs and eQTLs, as well as between pQTLs and complex traits were carried out using coloc (version 3.2.1) R package [50], which assumes that each locus has a single causal variant. Priors used for the colocalisation analysis were *P₁* = 10^−4^, *P₂* = 10^−4^ and *P₁₂* = 5×10^−6^, as suggested by Wallace et al. [51]. For each protein-level genome-wide–significant (*P* < 5 × 10^−8^) pQTL locus, we extracted regions in a 1-Mb radius of its lead variant to test for colocalisation. The results were considered significant when the posterior probability for colocalisation (PP4) exceeded 0.8.

In an pQTL–eQTL colocalisation analysis, we compared our significant pQTL loci to all eQTL Catalogue datasets [18], excluding those of Kasela et al. [52] and Lepik et al. [29] due to sample overlap, containing gene expression, exon expression, transcript usage and txrevise event usage data, and GTEx (version 8) [53] datasets containing gene expression data (https://www.ebi.ac.uk/eqtl/Methods/; S3 Table). We lifted the pQTL summary statistics over to an hg38 build to match with the eQTL Catalogue.

The region-wide associations for GWAS traits enrolled into the colocalisation analyses were extracted from the MRC IEU OpenGWAS database and were examined using the ieugwasr (version 0.1.5) R package (https://github.com/MRCIEU/ieugwasr; S4 Table). Since proteins were selected based on associated traits from the PheWAS, they were all associated with clinical traits (i.e. drugs, surgeries, diseases/conditions). In addition, all selected proteins except IL6R had primary pQTLs that did not include nonsynonymous variants, to minimise the possibility of association due to the epitope effect. IL6R was selected because it has been widely reported by previous pQTL studies as an example of the successful linking of molecular traits and diseases to discover drug targets [45,54]. The input data consisted of region-based summary statistics for six protein traits and 61 complex clinical traits.

### Two-sample MR

We conducted a two-sample MR analysis using protein levels with significant colocalisation (PP₄ ≥ 0.8) as exposures and complex traits as outcomes, using the TwoSampleMR (version 0.5.6) R package [55,56]. Independent variants obtained previously by clumping served as instrumental variables. We conducted the analysis using an inverse variance weighted fixed-effects method and a single instrument–based Wald ratio test. To correct for multiple testing, we adjusted *P-*values using false discovery rate (FDR) correction; results were considered significant at Benjamini-Hochberg FDR ≤ 0.05.

### CNV pQTLs, eQTLs and colocalisation

To determine whether any of the examined proteins are genetically regulated by larger structural variants, we conducted a pQTL mapping using CNV data. Description of the used CNV data is in the Methods section for CNV detection and quality control. Associations between previously described standardised protein measure residuals and CNV sites were assessed by using the MatrixeQTL R package [57]. The post–quality control sample sizes for the Inflammation, Oncology II, CVD II and CVD III panels were 481, 480, 489 and 488 unrelated (PI_HAT < 0.2) individuals, respectively. To discard rare CNV events, all CNV sites with in-sample frequencies of the most frequent copy number >0.95 were excluded. Additionally, unique non-overlapping CNVs were included. The final set used in the pQTL analyses comprised of 2,465 CNV sites [1,375 deletions (CN < 2), 482 duplications (CN > 2) and 608 combined deletions and duplications]. The genome-wide significance threshold was set to 1.12 × 10^−7^ (0.05 / 2,465 / 181).

For each significantly associated CNV, all SNP markers within a 500-kbp proximity were tested for potential tagging effects. For this purpose, the SNP pQTL analysis using EPACTS was repeated for these regions with the CNVs included as covariates.

The same CNVs were tested against the expression levels of 12,619 genes [29], and the CNV pQTL results were then cross-referenced with eQTLs identified from the same set of individuals. The eQTL results were corrected for multiple testing and a Bonferroni-corrected threshold of *P* < 1.61 × 10^−9^ [0.05 / (2,465 CNVs × 12,619 genes)] was applied. Overlapping eQTL–pQTL pairings were tested in an MR framework using the summary statistics–based ratio estimate (Wald test) [58], and Spearman’s rank correlation coefficient was calculated for gene expression vs protein expression in the same individuals. We hypothesised that CNVs in gene regions would be considerably more likely than other causal variants to modulate the expression of those genes; thus, non-zero ratio estimates were taken to indicate shared causal CNVs of gene expression and protein traits.

### PheWAS of CNV pQTLs

CNV pQTLs from primary mapping that reached genome-wide significance (*P* < 1.12 × 10^−7^) or the suggestive significance threshold (P < 2 × 10^−5^) were included in a PheWAS, resulting in the inclusion of 38 CNV regions. All data included in the PheWAS were obtained using the *lm* function with custom R scripts from 2,115 unrelated Estonian Genome Centre samples for which WGS data were available, and were corrected for age, sex and six genotype principal components (PCs; calculated from common SNPs). The 744 phenotypes examined were anthropometric traits (height, weight, body mass index, hip circumference, waist circumference, waist–hip circumferences ratio), cell counts from RNA-sequencing data (white blood cells, red blood cells, platelets, neutrophils, monocytes, lymphocytes, eosinophils, basophils), nuclear magnetic resonance spectroscopy–detected metabolites (*n* = 225) and International Classification of Diseases, 10th revision (ICD-10) diagnoses with at least 20 carriers in the sample (*n* = 505). Self-reported diagnoses not reported elsewhere were set to not available. Sex-specific diagnoses (ICD-10 codes F52, N4* and N5* for men, D25, D26, D27, E28, N7*, N8*, N9*, O* and Z3* for women) were analysed using only samples of the relevant sex as controls. The PheWAS significance threshold was set to *P* < 0.05 / 420, as 420 PCs calculated on all included phenotypes explained 95% of the variability.

### Identification of CNV-tagging SNPs for pQTLs

To aid the interpretation of the CNV-pQTL results, we examined additional pQTLs not detected in this study due to the small sample size or the lack of protein measurements, by using a CNV-tagging proxy SNP approach. To detect additional CNV–protein associations, we extracted all SNPs with MAFs > 0.01 from each common (major allele frequency < 0.95) CNV and its 500-kb flanking region, as identified in 2,230 Estonian WGS samples. We calculated Pearson correlation coefficients between the CNVs and SNPs using custom R scripts. SNPs with *R^2^* > 0.8 were defined as CNV-tagging proxy SNPs. The proxy SNPs were then compared with a published set of SNP pQTLs in two larger sets of unique proteins [4,9] to determine the degree of overlap. We used data on 1,021 independent autosomal lead pQTL variants for 1,478 proteins from the large-scale pQTL study conducted by Sun et al. [4]; 824 (80.7%) of these variants were present in the EstBB WGS dataset. We extended the analysis to include data from the largest pQTL study to date, conducted with 35,571 samples and resulting in the detection of 10,248 independent autosomal pQTLs for 1,463 proteins [9]. The two studies encompassed 2,438 unique proteins, enabling broader investigation. The resulting loci were reported as potential cases in which the underlying CNVs might be the causal variants. Figure depicting tagged-CNV pQTLs was done by using the RIdeogram v02.2.2 R package [59].

## Results

### Discovery of pQTLs

We identified 278 (184 *cis* and 94 *trans*) pQTLs for 157 (48.2%) of the 326 proteins examined, using a protein-level genome-wide significance threshold of *P* < 5 × 10^−8^ (S2 Table). When using a strict multiple testing correction threshold of *P* < 2.76 × 10^−10^, 151 pQTLs (131 *cis* and 20 *trans*) for 99 proteins remained significant (S2 Table). All interpretative analyses were conducted using protein-level genome-wide-significant results.

To provide a comparison with previous research, we compared our results with previously published data. From the Pietzner et al. study [7], 147 pQTLs (52.88%) were nominally significant (*P* < 0.05) and accessible for comparisons. After correcting for multiple testing, 147 pQTLs remained significant (Benjamini-Hochberg FDR < 0.05) and 91.84% (135/147) of pQTLs were directionally concordant with the current study (S2 Table). 66.19% (184/278) of pQTLs were tested in the Sun et al. study [4]. Of them, 55.98% (103/184) were significant (Benjamini-Hochberg FDR < 0.05) and 89.32% (92/103) were directionally concordant (S2 Table). 7.55% (21/278) pQTLs were also tested in the Suhre et al. study [46] and 57.14% (12/21) were significant (Benjamini-Hochberg FDR < 0.05), and all the significant pQTLs were directionally concordant with the current study (S2 Table). 12.23% (34/278) pQTLs were tested in the Folkersen et al. study [45] and 85.29% (29/34) of the pQTLs were significant (Benjamini-Hochberg FDR < 0.05) and all the significant pQTLs were also directionally concordant with the current study (S2 Table). Concordance with previous studies demonstrates the robustness of our results.

Fourteen (4.3%) of the proteins were measured in multiple arrays. Associations for the CXCL1, CCL3 and VEGFA proteins were validated by multiple independent arrays, in which the same genetic regions reached genome-wide significance and showed concordant effect directions. The total numbers of associated proteins were similar for all panels and ranged from 38 to 43 (S2 Table). The detected associations included 278 independent pQTL variants [184 (66.2%) *cis* and 94 (33.8%) *trans*], 9.35% of which were indels. Of the 157 associated proteins, 61 (38.9%) had more than one independent pQTL. Twenty-one proteins had both *cis* and *trans* associations. A MICA-MICB heterodimer coded from the chromosome 6 *HLA* region had the largest number of independent associations (*n* = 12; Fig. 2A). In concordance with previous studies [4,9,60], there was an inverse relationship between the effect size and MAF (Fig. 2B), and the associations were the strongest for significant *cis*-pQTL variants located nearest to the TSSs of the relevant protein genes (Fig. 2C). The largest proportion of these *cis*-pQTLs [*n* = 73 (39.7%)] was located in intronic regions (Fig. 2D). Of the 184 *cis* associations detected for 104 proteins, 31 (16.85%) were with protein-altering primary lead *cis*-pQTL variants and an additional 5 were with *cis*-pQTL variants in high LD with PAVs. These 36 (12.5%) pQTLs were designated as potential pseudo-pQTLs because currently it is difficult to exclude the possibility of technical signal happening due to the difference in antibody binding affinity.

**Figure 2.**
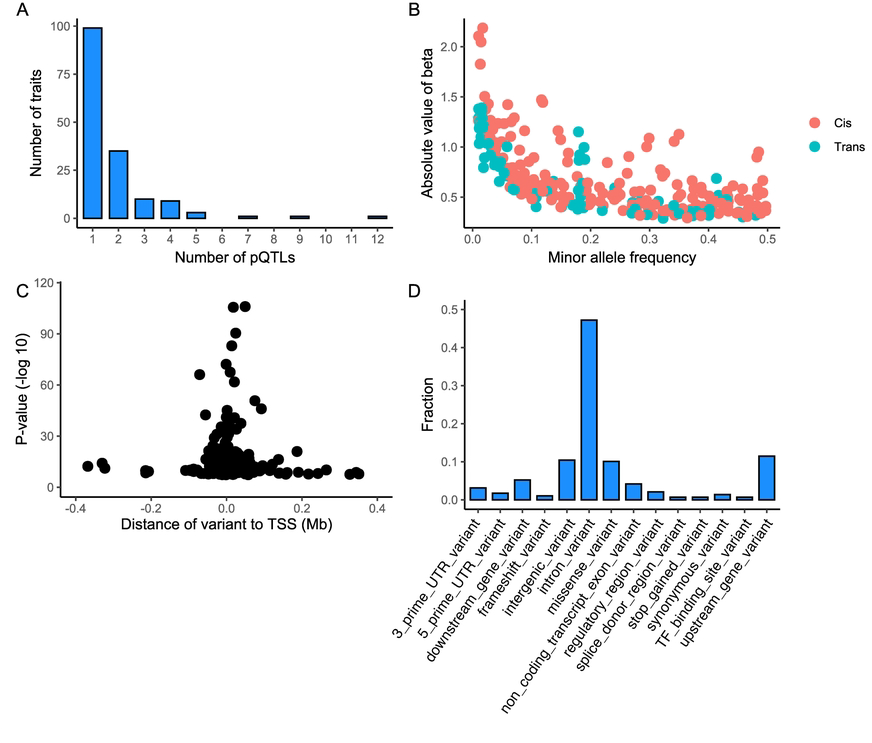
A. Numbers of genome-wide significant associations of variants with protein traits. B. Absolute beta values according to minor allele frequencies (MAFs). C. Significance of primary pQTL mapping cis associations according to distances from transcription start sites (TSSs). D. Functional annotation classes for the top cis variants from pQTL mapping, expressed as fractions.

The strongest *cis* association was between the missense variant rs2228145 (p.Asp358Ala) and the IL6RA level (MAF = 0.35, *P* = 1.04 × 10^−106^). Additional strong *cis* associations included the rs1569960 and the SIRPA level (MAF = 0.34, *P* = 2.67 × 10^−106^) association, with four independent signals in the SIRPA *cis* region; and a frameshift-causing insertion rs139130389 and the FOLR3 level (MAF = 0.12, *P* = 3.91 × 10^−91^) association, with three independent signals in the FOLR3 *cis* region.

The most significant *trans* association was that of the *PLAUR* missense variant rs4760, located on chromosome 19, affecting the level of TNFRSF10C (8p21.3; MAF = 0.18, *P* = 4.60 × 10^−56^). Strong *trans* associations were between the rs8176671 and the CDH5 level (16q21; MAF = 0.19, *P* = 8.83 × 10^−40^) as well as between the deletion rs8176643 and the SELE level (1q24.2; MAF = 0.18, *P* = 7.98 × 10^−36^); both of these variants are intronic variants for the 9q34.2 locus of the *ABO* gene. This locus was a *trans*-signal hotspot, with intronic variants additionally associated with the ICAM2, galectin-4 (LGALS4), PODXL and LIFR protein levels. Additional *ABO* variant rs12216891 was associated with the CTRC level (MAF = 0.19, *P* = 8.39×10^−30^).

Two of the proteins examined (MICA/B and IL27) are heterodimers, made up of multiple subunits that are translated from two different genes at distinct loci. For IL27, we identified one independent *trans* signal for an intronic variant for CCDC94 (rs56075200; MAF = 0.32, *P* = 8.62 × 10^−35^). For MICA/B, we identified ten independent signals in the *cis* region of one subunit on chromosome 6 (the strongest signal was for an intronic variant of MICA: rs3132467; MAF = 0.30, *P* = 3.04 × 10^−68^) and two *trans* associations.

To determine if there were any corresponding eQTLs for pQTLs, we conducted an eQTL analysis, using the whole blood gene expression data from the same individuals and the same time point. Gene expression data was available for 109 proteins with 201 pQTLs, including two heterodimers with two subunits encoding the protein. In total, we detected 62 significant (Benjamini-Hochberg FDR < 0.05) eQTLs (59 *cis*, 3 *trans*) (S5 Table). 77% (48/62) of them were directionally concordant with corresponding pQTLs.

We found that 95% CSs for 151 proteins were linked to 131 independent genomic loci (S6 Table). LDLR, TNFRSF11B, TNFRSF6B, WISP1, CXCL1 and PLAU proteins showed significant pQTL effects but yielded no CS. Signals for CCL3, CXCL1 and VEGFA from multiple assays were also validated by fine mapping to the same genetic regions. The 95% CSs contained an average of 15.7 variants (*cis* sets, 15.76; *trans* sets, 15.6). Fifty-five (36.4%) proteins had single-variant CSs. Of the 31 proteins with single-variant CSs in *cis* regions, 13 were fine-mapped to lead PAVs from primary pQTL mapping. Thirty-three (32.7%) out of 101 *cis* regions were fine-mapped to more than one signal (mean, 1.4 signals/region), with the CCL24 *cis* region having the largest number of independent CSs (*n* = 5). In contrast, all associated regions for pQTL *trans* signals were fine-mapped to a single CS.

Since a large proportion (217/278) of primary pQTLs were located in intergenic and intronic regions, we queried RegulomeDB [32] to establish the variants’ potential regulatory function. We obtained regulatory information for 260 of 278 pQTLs corresponding to 251 unique lead variants. Eleven variants (all *cis*) were previously established eQTLs and had evidence for transcription factor binding– and/or DNase peak–related functions. Seventeen lead variants (12 *cis* and 5 *trans*) had chromatin immunoprecipitation sequencing– and DNase-based evidence for regulatory functions, but were not eQTLs (S7 Table).

### pQTL–eQTL colocalisation

The pQTL–eQTL colocalisation analysis was performed with 198 pQTL loci (corresponding to 157 unique proteins), 18 eQTL Catalogue datasets and GTEx tissue eQTL data. We identified 14,064 cases of pQTL–eQTL colocalisation (PP4>0.8), involving 105 proteins [7,936 (56.4%) *cis*- and 6,128 (43.6%) *trans*-pQTLs; Tables 1, S8]. Colocalisations classified as *cis* consisted of 2,021 (25.5%) cases in which colocalising eQTLs and pQTLs affected the same gene product and 5,915 (74.5%) cases in which the colocalising loci affected different gene products in the *cis* regions. *Cis* and *trans* pairs were specific to 73 and 26 proteins, respectively, and 6 proteins (IL1R2, TEK, MIA, FCRLB, PDCD1LG2 and MICA-MICB) had colocalisations for both *cis* and *trans* associations. The largest number of colocalisations was found for pQTLs of the MICA-MICB heterodimer (*n* = 6,583), followed by OSCAR (*n* = 1,207) and ACP5 (*n* = 1,105) pQTLs.

**Table 1.**
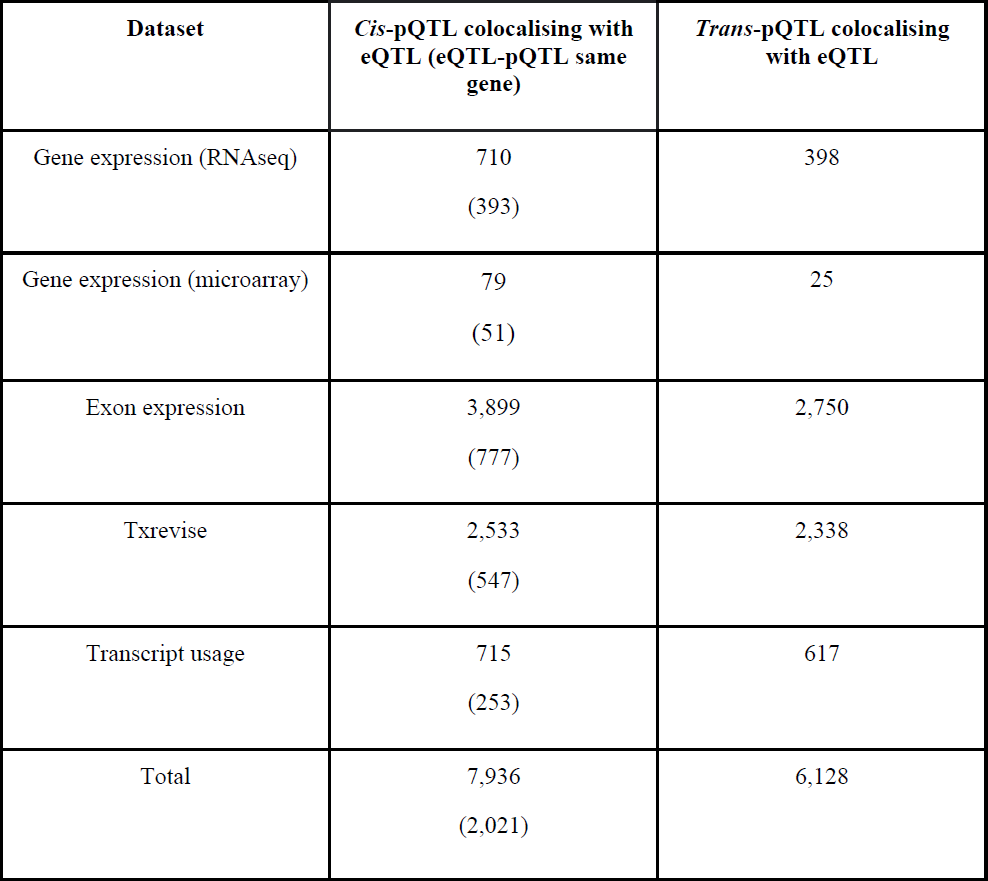
Overview of significant colocalisation events for eQTLs from eQTL Catalogue datasets and pQTLs. The numbers of colocalisation with genes encoding corresponding proteins are shown in parentheses.

| Dataset | *Cis*-pQTL colocalising with eQTL (eQTL-pQTL same gene) | *Trans*-pQTL colocalising with eQTL |
| --- | --- | --- |
| Gene expression (RNAseq) | 710<br>(393) | 398 |
| Gene expression (microarray) | 79<br>(51) | 25 |
| Exon expression | 3,899<br>(777) | 2,750 |
| Txrevise | 2,533<br>(547) | 2,338 |
| Transcript usage | 715<br>(253) | 617 |
| Total | 7,936<br>(2,021) | 6,128 |

Since the protein measurements originated from blood, the most widely studied tissue, the largest fraction of pQTLs colocalised with blood eQTLs. However, while using the GTEx dataset, we also found 739 cases of pQTL–eQTL colocalisation in multiple tissues (Fig. 3, S8 Table). For 55 proteins with *cis*-pQTLs, 503 (68.1%) colocalising eQTLs were identified; for 22 proteins with *trans*-pQTLs, 236 (31.9%) colocalising eQTLs were identified. *Cis*-pQTLs colocalising with eQTLs were detected in 49 tissues, and *trans*-pQTLs colocalising with eQTLs were identified in 46 tissues (not in Epstein-Barr virus–transformed lymphocytes or uterine or vaginal tissue).

**Figure 3.**
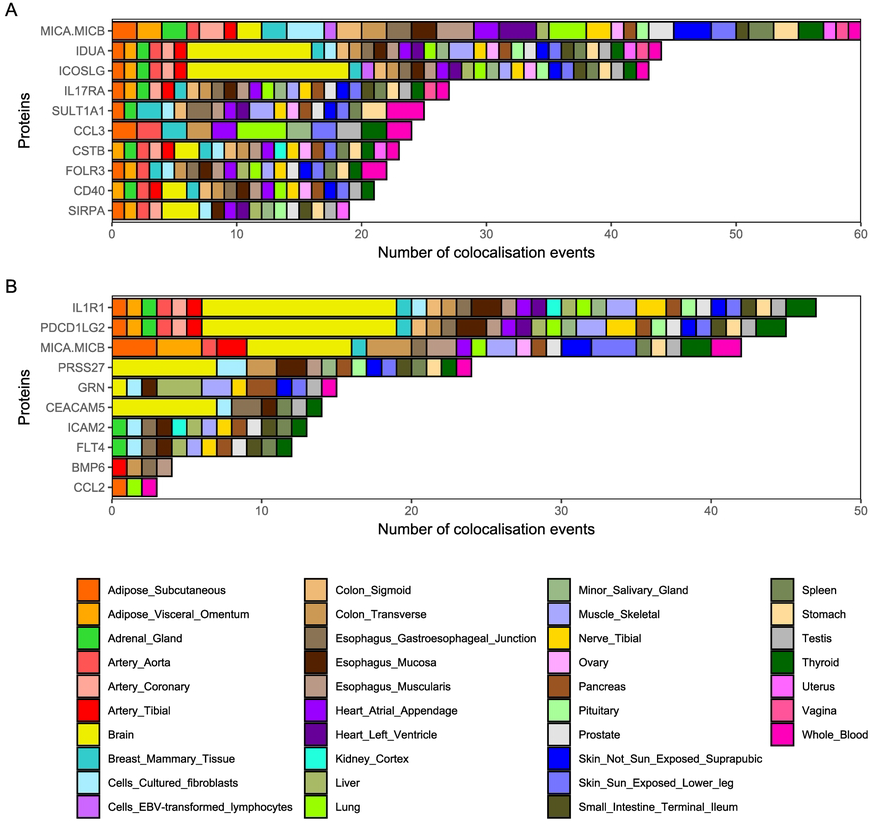
Overview of 10 *cis*-pQTL (A) and *trans*-pQTL (B) proteins with the most colocalising eQTLs from the GTEx database (version 8; GTEx Consortium, 2020) Colours indicate eQTL tissues of origin. Brain tissues are pooled; a complete list is provided in S8 Table.

### PheWAS on metabolite and epigenetic QTLs

Queries for the 268 lead pQTL variants led to the identification of 17 variants (from 6 *cis* and 13 *trans* associations for 18 proteins) associated with 160 metabolite traits (S9 Table). The majority [*n* = 158 (52.3%)] of the mQTLs discovered were for the *APOE* missense variant rs7412, which had a *trans* association with the level of LDLR. Four metabolic traits [apolipoprotein B, the concentration of very small very-low-density lipoprotein (VLDL) particles, and phospholipids and total lipids in very small VLDL] had seven associations each.

From the epigenetic QTL datasets, we identified 6,236 meQTLs, 267 histone modification QTLs and 129 exon-inclusion PSI associations for 193 primary pQTLs (from 142 *cis* and 60 *trans* associations for 130 proteins; S10 Table). Most (*n* = 256) meQTLs were associated with the ADAM8 *cis*-pQTL rs2995310. The variant with the most (*n* = 10) histone modifications was rs10415777, a *cis*-pQTL for OSCAR. Methylation data originates from five tissues: cord blood, monocytes, neutrophils, T cells and whole blood; due to tissue availability, 78.7% (4,906/6,236) of the identified meQTLs were from whole blood studies.

### Common SNP pQTLs and complex traits

#### PheWAS

The queries for the 268 unique lead variants and high-LD proxies led to the identification of 135 (50.4%) variants with 5,046 significant associations for 432 complex traits (S11 Table). Of these associations, 1583 (31.4%) were with various blood cell traits from the study conducted by Astle et al. [61]. As expected, given the targeted nature of our protein panels, coronary artery disease (CAD) and rheumatoid arthritis were most often linked to pQTLs with 118 and 99 associations, respectively. For example, 5 of 145 significant independent signals for CAD from mixed-ancestry samples [62] and 2 of 7 significant loci for rheumatoid arthritis from the study conducted by Stahl et al. [63] were pQTLs in our dataset. In terms of the most associations per pQTL lead variant, *ABO* intronic variant rs507666 had the most associations per lead pQTL variant [*n* = 332, 85 (25.6%) with blood cell traits]. No associated traits were found for 62 proteins.

For 61 proteins (64 lead pQTL variants, 36 *cis*- and 28 *trans*-pQTLs), significant associations were detected in both the eQTL colocalisation analysis and PheWAS. We restricted this set to 27 proteins (28 variants) which were not coded from the *HLA* region but showed associations with diagnosis, treatment, or other phenotypes linked directly to health status (excluding haematological and biochemical measurements). Six of these proteins (CD6, PRSS27, CEACAM5, CD40, TNFRSF6B and IL1RL1) had significant colocalisations with eQTLs from brain tissue, but no evidence of shared conditions with direct effects on the brain tissue in the PheWAS.

For example, based on pQTL-eQTL colocalisation analysis, IL6R pQTL signal was also an eQTL of the *IL6R* gene in macrophages, monocytes, T cells, whole blood and pancreatic islets. A previous study has shown a link between IL6R and CAD [64]. We also identified associations between IL6R pQTLs and CAD, rheumatoid arthritis and 7 other disease traits (S11 Table), thereby supporting the findings of the study [63]. As another example, *IL1RL1* pQTLs colocalised with *IL1RL1*, *IL18R1* and *IL18RAP* eQTLs detected in multiple cell types with direct effects on the immune system (e.g. T-cells; S8 Table); these variants were associated with asthma and allergic reactions in the PheWAS.

Eleven out of 27 proteins had *trans*-associations. *Trans*-pQTLs for the CTRC and TEK proteins were in the *ABO* locus and colocalised with *ABO* eQTLs; in the PheWAS, they were linked to multiple self-reported diagnoses (e.g. ‘blood clot in the leg’) from the UK Biobank sample, and to haematological traits.

Most [*n* = 140/157 (89.2%)] of the proteins with significant pQTLs belonged to the druggable genome category. These proteins were associated with 1,365 drug–gene interactions.

#### Colocalisation analysis

Based on the pQTL associations with genetic regions, PheWAS and eQTL colocalisation results, we chose five *cis*-pQTL effects (affecting FGF5, IL1RL2, TNFRSF6B, IL2RA, and IL6R) that were associated with clinical traits and had significant pQTL-eQTL colocalisations. Furthermore, SULT1A1 was chosen due to additional CNV–pQTL associations in its region which enabled to analyse colocalisation with respective complex traits. All selected proteins except IL6R had synonymous lead pQTL variants. Therefore, the input data for colocalisation analyses comprised of region-based summary statistics for 6 protein traits and 61 clinical complex traits (83 pQTL–complex trait pairs).

We identified 46 significant colocalisation events (S12 Table). FGF5 had 25 colocalisations with cardiovascular phenotypes and medications, such as CAD and perindopril use. IL6R had a total of 11 significant colocalisations, which included colocalisations with CAD as well as immunological conditions such as asthma. TNFRSF6B and SULT1A1 colocalised with inflammatory bowel disease, and TNFRSF6B also separately colocalised with its two main forms: Crohn’s disease and ulcerative colitis. IL2RA colocalised with tonsillectomy +/− adenoid operation. The PheWAS revealed associations of IL1RL2 with immune diseases which were not supported by the colocalisation results.

#### MR findings

We conducted MR analyses using 46 significant (FDR-corrected) pQTL–complex trait pairs from the colocalisation analysis (Fig. 4, S13 Table). We found a causal relationship between the elevated level of soluble IL6R and a lower risk of cardiovascular disease (*P* = 2.35 × 10^−24^, Benjamini-Hochberg FDR = 1.08 × 10^−22^). Higher IL6R levels were also associated with an increased risk of inflammatory conditions such as asthma and eczema (*P* = 2.04 × 10^−4^, Benjamini-Hochberg FDR = 2.60 × 10^−4^; *P* = 1.24 × 10^−5^, Benjamini-Hochberg FDR = 1.96 × 10^−5^, respectively). The TNFRSF6B level was causally linked to a reduced risk of inflammatory bowel disease and its subtypes (inflammatory bowel disease (A294), *P* = 4.00 × 10^−20^, Benjamini-Hochberg FDR = 9.19 × 10^−19^; Crohn’s disease (A12), *P* = 1.18 × 10^−16^, Benjamini-Hochberg FDR = 1.82 × 10^−15^; ulcerative colitis (A970), *P* = 2.14 × 10^−8^, Benjamini-Hochberg FDR = 7.56 × 10^−8^). Elevated levels of FGF5 were associated with a significantly increased risk of coronary disease (*P* = 8.94 × 10^−6^ and Benjamini-Hochberg FDR = 1.47 × 10^−5^).

**Figure 4.**
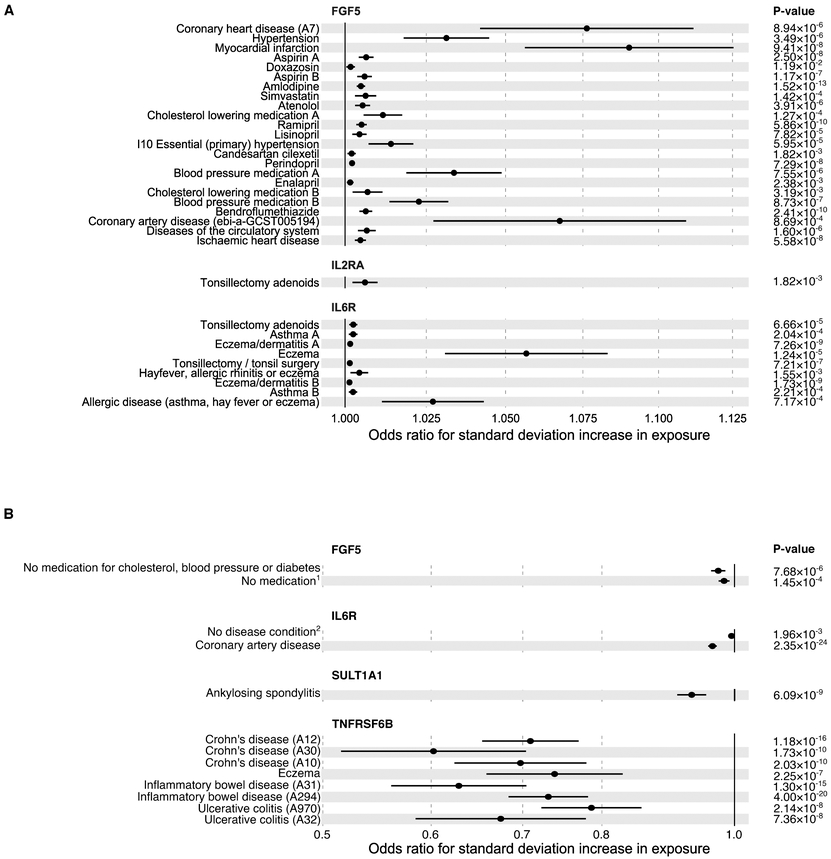
Forest plots of Mendelian randomisation results for proteins with positive (A) and negative (B) effects on complex traits. Protein (exposure) names are indicated on top of the section, complex traits (outcomes) are on the left side. Multiple instances of traits with the same name for one protein, indicating MR signal replication across multiple studies of the same trait, have been marked ‘A’ and ‘B’. Error bars denote standard errors and all presented results are significant at a Benjamini-Hochberg FDR < 0.05. Details of causal associations are provided in S13 Table. ^1^“Medication for cholesterol, blood pressure, diabetes, or take exogenous hormones: None of the above” (MRC IEU UK Biobank); ^2^“Blood clot, DVT, bronchitis, emphysema, asthma, rhinitis, eczema, allergy diagnosed by doctor: None of the above” (MRC IEU UK Biobank).

### Rare variant pQTLs

The gene-based association analysis revealed 19 significant associations [5 (26.3%) *cis* and 14 (73.7%) *trans*] emanating from 19 genes containing rare nonsynonymous SNPs and affecting the levels of 7 proteins (S14 Table). The majority of identified rare variant effects (13, 68.4%) were with the level of GDF-15. The most significant rare variant association was a *trans* signal between *JAKMIP1* on chromosome 4 and the level of GDF-15 (*P* = 5.41 × 10^−12^). We also assessed if rare nonsynonymous SNPs affect the expression of same genes encoding the corresponding pQTL proteins, however we did not detect any nominally significant (Benjamini-Hochberg FDR < 0.05) associations (S14 Table).

We next conducted GeneMANIA network analysis [37, 38] to identify functional connections between genes harbouring rare SNPs and proteins affected by *trans* associations. First, we studied the potential connection between rare variant genes associated with the GDF-15 level. Ten of the identified genes harbouring rare SNPs (*CKAP5*, *GDF15*, *JAKMIP1*, *KRT19*, *STAT5B*, *SLC35E1*, *RNF112*, *TUBGCP4*, *ZNF766* and *PPAPDC1B*), including gene encoding identified pQTL protein, formed shared network with GDF-15, based on co-expression (57.85%), pathway (19.97%), physical (18.45%) and genetic (3.73%) interactions, according to GeneMANIA. However, no functional connection to GDF-15 was found for *LY6G6E*, *RPL7L1* and *EFR3B*. *Trans* associations between rare variants and SELPLG and MUC-16 levels were supported by the GeneMANIA-based identification of two shared networks: between *TMEM119* and SELPLG, as well as *GAL3ST2* and MUC-16, respectively. Those connections were based mainly on physical interactions (77.64%) and co-expression (8.01%).

Four proteins (CTSZ, GDF-15, PON3 and SELPLG), had significant associations from both, common variant and rare variant pQTL analyses. For CTSZ and GDF-15, the genetic regions detected from the rare variant analysis were not the same as identified by SNP pQTL analysis. However, PON3 had direct *cis* associations emanating from from 7q21.3 locus in both analyses: nonsynonymous variants of *PON3* in the rare variant pQTL analysis and rs10953142 in the common variant pQTL analysis. Similarly, SELPLG had *cis* associations emanating from 12q24.11 locus: nonsynonymous variants of the *TMEM119* for rare variant analysis and an intergenic rs11114010 for common variant analysis.

### CNV pQTLs

We detected 12 significant (Bonferroni-corrected *P*-value threshold 1.12 × 10^−7^) pQTL associations between CNVs and plasma protein levels (7 *cis* and 5 *trans*, 11 proteins; S15 Table), with two *cis* associations detected for the MICA-MICB heterodimer. The CNV eQTL analysis in the overlapping set of samples identified 673 significant (Bonferroni-corrected *P*-value threshold 1.61 × 10^−9^) CNV eQTLs for 244 unique genes (S16 Table). 16.67% (2/12) of significant CNV pQTLs had significant CNV eQTL associations with a corresponding gene.

For example, the deletion in the 3q12.1 intergenic region (chromosome 3: 98,410,653-98,414,807 bp; frequency = 0.651) acted as a hub, having multiple *trans* associations with protein levels: ICAM2 (*P* = 1.31 × 10^−29^), FLT4 (*P* = 2.34 × 10^−24^), PDCD1LG2 (*P* = 2.88 × 10^−15^) and IL1R1 (*P* = 8.19 × 10^−8^). Three of these associations (with ICAM2, FLT4 and PDCD1LG2) were also detected by the SNP pQTL analysis but did not remain significant after conditioning of the model on the CNVs, suggesting that CNV may underlie the observed associations. However, eQTL analysis indicated that none of the genes encoding those proteins is regulated by this locus, and a follow-up GeneMANIA network analysis [37,38] revealed a shared network based on physical interactions (77.64%), co-expression (8.01%), predicted functional relationship between genes (5.37%), co-localisation (3.63%), genetic interactions (2.87%), pathway (1.88%) and shared protein domains (0.60%).

Another *trans* association example was between a 5q13.2 CNV (chromosome 5: 70,305,253– 70,312,310 bp; deletion frequency = 0.074, duplication frequency = 0.195) overlapping the *NAIP* gene but affecting IL-18 level (*P* = 7.9 × 10^−10^). This locus was also an eQTL for *NAIP* (*P* = 6.4 × 10^−48^), but not for IL18 expression (*P* > 0.001). We also detected moderate correlation between IL-18 protein expression and *NAIP* gene expression (Spearman’s *R* = 0.17); Spearman correlation coefficient between IL-18 protein and gene expressions was 0.05. MR analysis using *NAIP* gene expression as exposure and IL-18 level as an outcome confirmed the causal effect of the CNV on the IL-18 protein level (Wald test; *Z* = 6.26, *P* = 3.8 × 10^−10^). This association was not observed in the SNP-based analyses, highlighting the case where the pQTL signal would not be detected.

From *cis* effects, we detected an association between CNV in the 16p11.2 region (deletion frequency = 0.022, duplication frequency = 0.382; partially overlapping SULT1A1; pQTL, *P* = 3.46 × 10^−21^; eQTL, *P* = 4.74 × 10^−119^) and SULT1A1 protein and gene expression. Similarly, we determined that a 19q13.42 deletion (frequency = 0.291) overlapping the *VSTM1* gene was an eQTL and a pQTL for nearby gene *OSCAR* (*P* = 1.77 × 10^−14^ and *P* = 5.64 × 10^−9^, respectively). However, the CNV was also associated with the expression of *VSTM1* itself (*P* = 1.81 × 10⁻^39^) and both gene–protein expression pairs showed moderate correlation (*OSCAR–* OSCAR, Spearman’s *R* = 0.32; *VSTM1–*OSCAR, Spearman’s *R* = 0.34). The effect of the CNV through gene expression is supported by the MR analysis, when using a CNV as an instrument, *OSCAR* expression as an exposure and OSCAR level as an outcome (*Z* = 5.94; *P* = 2.81 × 10^−9^) and secondly, VSTM1 as an exposure and OSCAR level as an outcome (*Z* = 5.92; *P* = 3.27 × 10^−9^). Those results suggest that CNV works through gene expression, although it remains unclear whether the effect on the OSCAR level is through *OSCAR* or *VSTM1* gene expression.

Additionally, we identified an association between the SIRPA level and a high-frequency (frequency = 0.955) 20p13 deletion overlapping *SIRPB1*, a paralog of *SIRPA* (*P* = 1.4 × 10^−11^; Fig. 5A and 5B). eQTL analysis indicated that the deletion was also associated with *SIRPB1*, but not *SIRPA*, expression (*P* = 3.5 × 10^−87^). The correlation between SIRPA protein and gene expression was weaker than that between SIRPA protein and *SIRPB1* expression (Spearman’s *R* = 0.075 and 0.202, respectively). Colocalisation was confirmed by the Wald test (*Z* = 6.92, *P* = 4.5 × 10^−12^; Fig. 5C). In SNP pQTL fine mapping, we detected two independent CSs, overlapping *SIRPB1* [variant with the largest posterior inclusion probability (PIP) = 0.295] and at *SIRPA* (variant with the largest PIP = 0.242; Fig. 5A). When conditioned on the deletion, the significance of pQTLs from only the *SIRPB1* CNV region was reduced dramatically (chr 20 position 1546911 variant pQTL mapping, *P*_primary_= 3.75 × 10^−11^, *P*_conditional_ = 0.41, regional pQTL mapping with EMMAX linear mixed-model [30] and the occurrence of the CNV and the number of its copies used as an additional covariate). This example highlights that the second signal from the primary pQTL analysis *SIRPA* locus was due to CNV-tagging variants rather than an independent signal.

**Figure 5.**
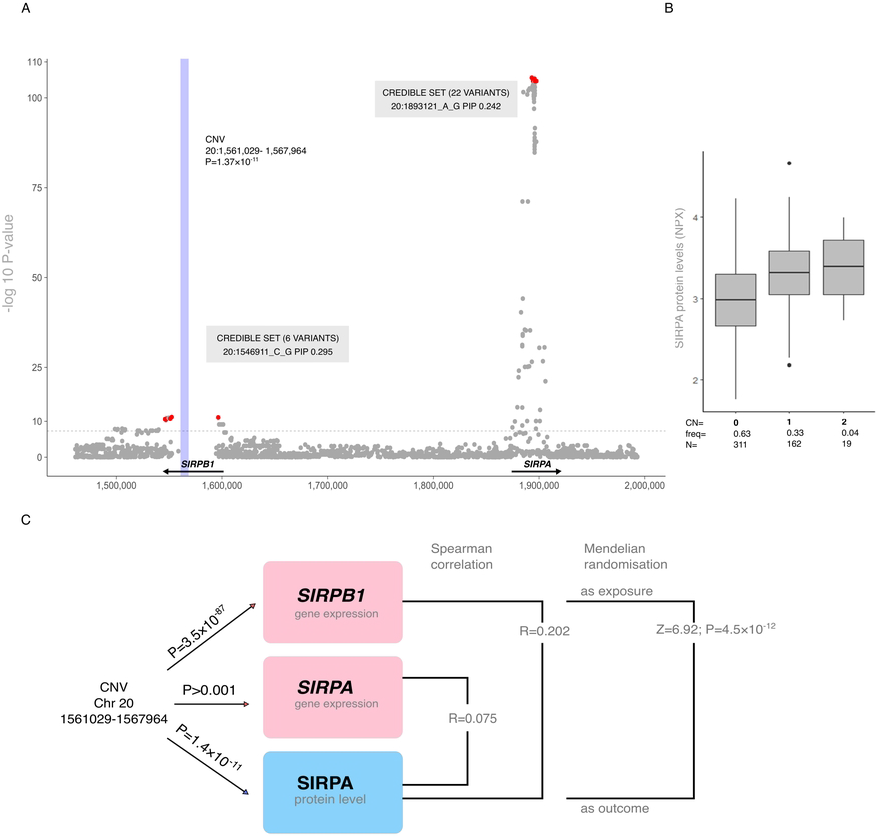
A. Regional plot combining SNP- and CNV-based results for the SIRPA level with additional single-variant fine-mapping information. The blue rectangle indicates the genetic location of the CNV. The horizontal dashed line indicates the genome-wide significance threshold of *P* = 5 × 10^−8^. Genetic variants identified by fine mapping as belonging to 95% credible sets are coloured red. The number of variants and the variant with the highest PIP in the credible set are indicated in grey boxes. B. Box plot of SIRPA levels based on the CNV number of copies and frequencies. Error bars indicate 95% confidence intervals; the bottoms and tops of the boxes are the 25th and 75th percentiles, respectively; the lines inside the boxes indicate medians. Outliers are depicted as circles. C. Overview of SIRPA level analyses. *P*-values are from the CNV-based pQTL analysis for SIRPA and eQTL analyses for *SIRPB1* and *SIRPA*.

Associations for nine proteins significant in both, CNV and pQTL mapping, were emanating from the same loci in both analyses. For example, ICAM2 and FLT4 had *trans* associations with rs12493830 on chromosome 3 and a CNV (chromosome 3: 98,410,653–98,414,807 bp) in the same intergenic region, separated by 3859 bp.

### PheWAS for CNV pQTLs

Significant PheWAS associations were detected for three CNVs. For the MICA-MICB dimer pQTL, associations were detected between CNV on chromosome 6 (31,292,078–31,293,977 bp; deletion frequency = 0.876) and medium HDL triglycerides (*P* = 8.82 × 10^−5^), and between a CNV on chromosome 6 (31,337,848–31,341,642 bp; deletion frequency = 0.074) and lower-limb oedema (ICD-10 code R60; *P* = 9.06 × 10^−5^). Additionally, we detected nominally significant associations for a CNV on chromosome 19 (41,381,588–41,387,347 bp, deletion frequency = 0.054 and duplication frequency = 0.022) with the pQTL of the MIA protein level (*P* = 2.38 × 10^−6^) and migraine (ICD-10 code G43; *P* = 3.14 × 10^−5^).

### CNV-tagging SNPs

To further interpret the of CNV-pQTL results, we examined additional pQTLs for proteins that were not measured in our study. For that, we leveraged LD between the EstBB CNVs and previously reported pQTL SNPs and prioritised CNVs which could underlie the previously reported pQTL associations (*R^2^* between SNP and CNV >0.8). We identified eight CNVs with possible effects on protein levels (Table 2) from the Sun et al. 2018 study [4]. Only one of those associations [proxy SNP rs10935473 with the CNV on chromosome 3 (98,410,653–98,414,807; deletion frequency = 0.651)] affecting FLT4/VEGF-sR3 levels, was identified in our study because the other proteins were not measured in our cohort.

**Table 2.**
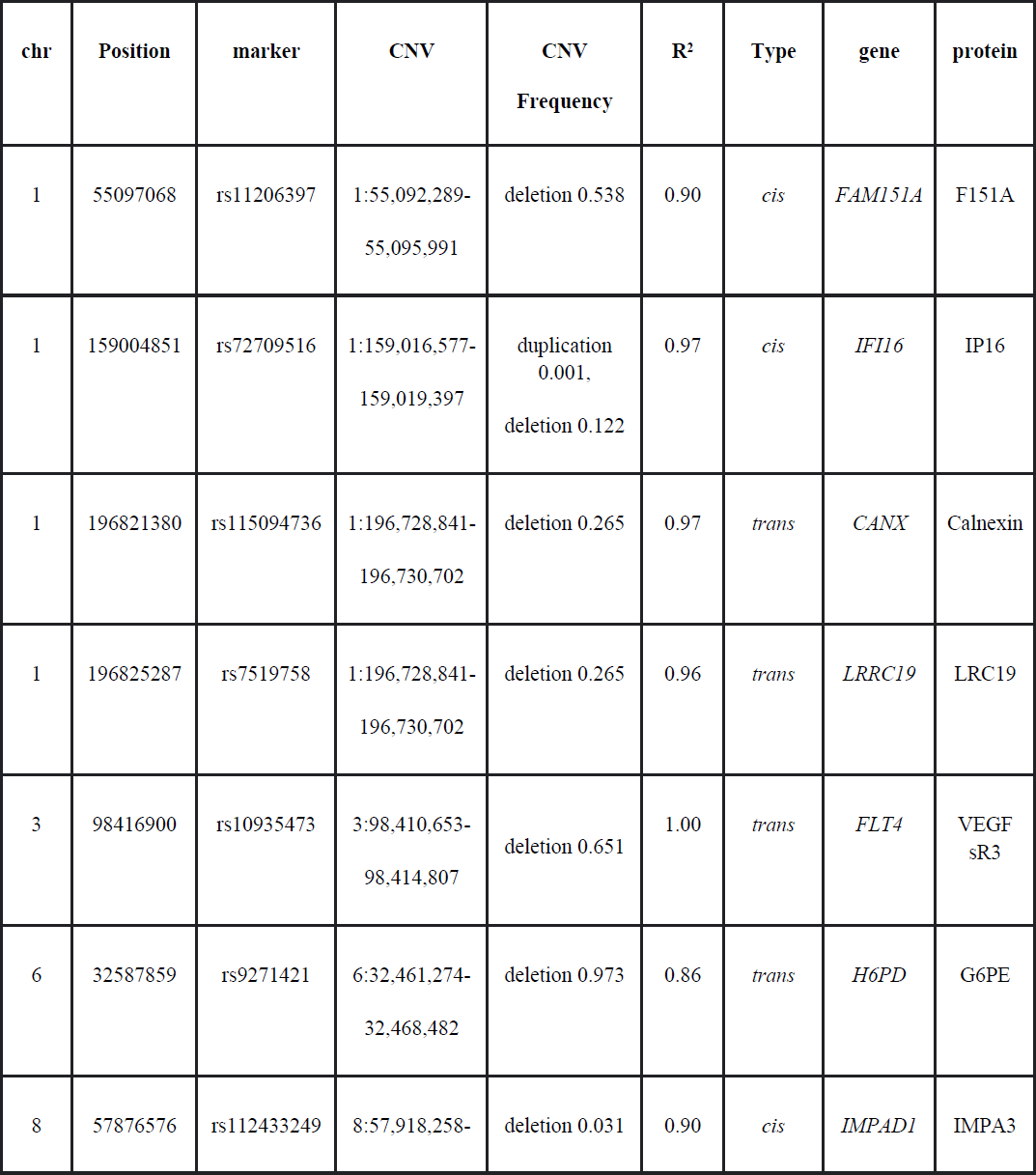
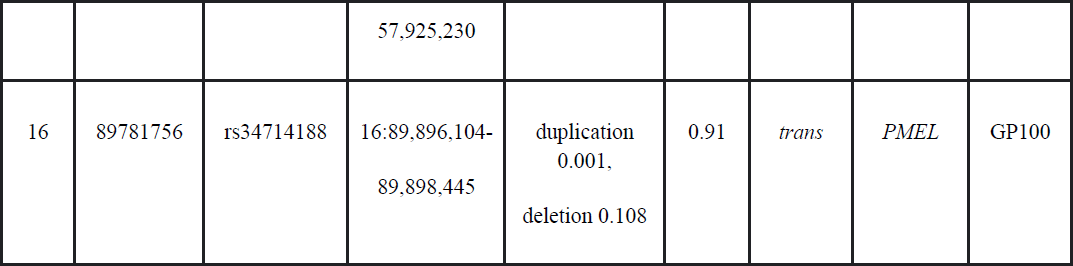
Overview of SNPs tagging CNVs for proteins reported by Sun et al. (2018) CNV frequencies are derived from the EstBB data.

We also detected 76 tagging SNP–CNV pairs for 33 unique CNVs and 72 proteins (S17 Table) from a more recent Sun et al. 2022 study [9]. Twenty-nine (40.3%) of the proteins were also measured in the EstBB cohort, of which six proteins had significant CNV pQTLs (*P* < 1.12 × 10^−7^). However, CNV-based pQTLs of the MICA-MICB heterodimer and SIRPA were not associated with the same CNVs in the EstBB cohort as tagged by SNPs in Sun et al.’s [9] study. Twenty-five (32.9%) of the tagging SNP–CNV pairs were associated with a deletion in the 3q12.1 intergenic region (chromosome 3: 98,410,653–98,414,807 bp, frequency = 0.651; the closest gene is *ST3GAL6*), a *trans* association hub (Fig. 6), and the same deletion was associated with four proteins (ICAM2, FLT4, PDCD1LG2 and IL1R1) in the EstBB dataset.

**Figure 6.**
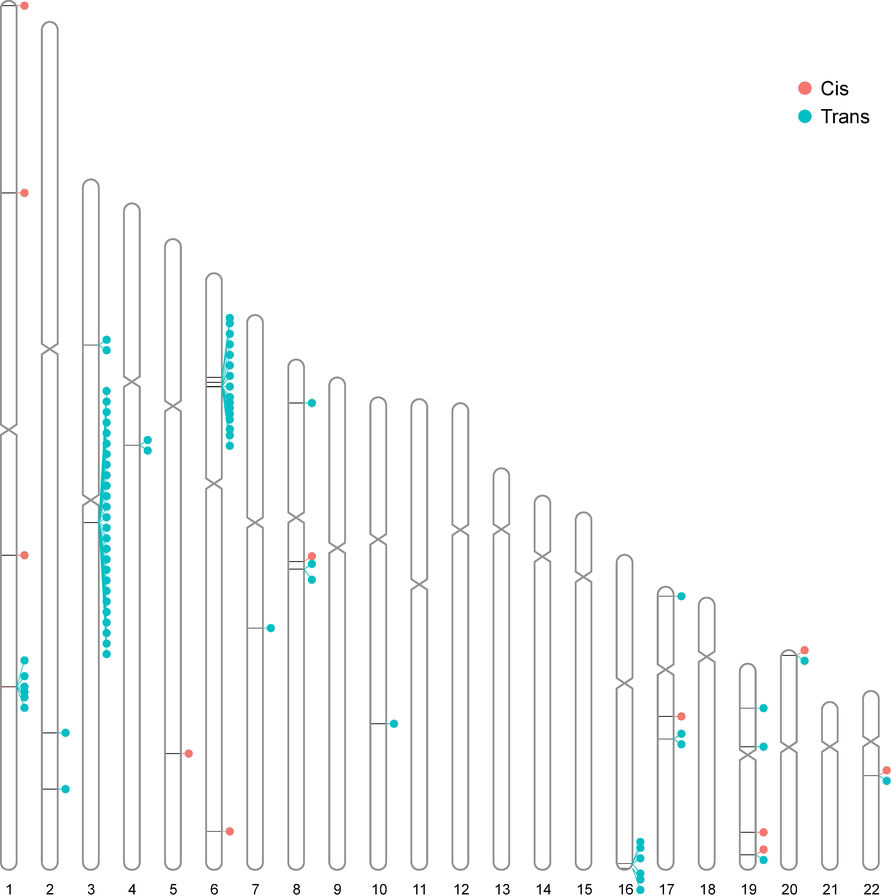
Overview of SNP-tagged CNV and protein *cis* and *trans* associations. Each line depicts the CNV which is in LD (*R^2^*>0.8) with pQTL SNP previously reported by Sun et al. (2018) or Sun et al. (2022) study. Each dot indicates corresponding pQTL protein and colour depicts the type of association.

None of the pQTLs tagging the CNV has known associations with complex traits which are not cell type or metabolite related, according to the GWAS Catalog. In addition, 19.7% (15/76) of the CNVs paired with tagging SNP were located in the *HLA* region on chromosome 6. The proteins TACSTD2, CLEC5A, IL15 and SIGLEC9 were affected by multiple *trans*-pQTL SNPs tagging CNVs. Whereas we detected a CNV associated with the SIRPB1 level on chromosome 20 (1,556,917–1,561,028 bp, deletion frequency = 0.336) and a deletion in the same locus overlapping *SIRPB1* and affecting the SIRPA level and (more strongly) *SIRPB1* gene expression, based on Sun et al. tagging-CNV analysis, the SIRPB1 protein level was associated with a different CNV than was its gene expression.

## Discussion

The SNV-pQTL analyses conducted in this study revealed 278 genetic variants (184 *cis* and 94 *trans*, including indels), that were associated with the levels of 157 unique proteins. Consistent with previous findings [4,6,8], the largest proportion of *cis*-pQTLs was located in intronic and intergenic regions. The analysis of individual-level WGS data together with in-sample LD information, enabled us to pinpoint the likely causal variants with a good resolution through statistical fine mapping. This mapping led to the identification of at least one 95% CS for each of 98 (53%) *cis* and 87 (47%) *trans* signals. For 16 *cis* and 28 *trans* associations, we identified 95% CSs consisting of the single most likely causal variants, which are good candidates for further functional studies. Notably, the prioritised variants for nine (56%) of the single-variant CSs for *cis*-pQTLs had protein-altering effects. This observation outlines that it is important to consider technical epitope effects in the cis-pQTL analyses [65]. However, the identification of PAVs demonstrates that fine mapping is also helpful for prioritising biologically causal variants, because PAVs are likely to have a direct, albeit technical, effect on protein levels. Only a limited number of pQTL studies have conducted fine mapping [9,66] as one of the post-GWAS analyses. We and Zhang et al. [66] detected CSs for 58 (59.2%) protein *cis* regions using data from cohorts of European ancestry, and Sun et al. [9] fine mapped CSs in 127 (67.6%) genetic regions for 117 proteins, matching our findings. The 95% CSs contained an average of 15.7 variants in our study and 22.7 variants (9.6 cis and 29.4 trans) in that of Sun et al. [9]. Our CSs for *cis* associations contained an average of 15.76 variants, whereas Zhang et al. used imputed genotyping data and reported an average of 21.29 variants [66]. Generally smaller credible sets might outline the added value of WGS data on fine mapping performance.

To support our findings with orthogonal data, we used the most comprehensive publicly available eQTL resource, the eQTL Catalogue [18], to conduct eQTL–pQTL colocalisation analyses. Detected colocalisations were 56.4% for *cis*- and 43.6% for *trans*-pQTLs. Of the *cis* associations, 25.5% (2,021/7,936) colocalised with the eQTLs for the corresponding protein-encoding gene from the full eQTL Catalogue, while for the GTEx dataset alone it was 54.3% (273/503). Given the use of eQTL data from different tissues, this analysis reflects how pQTLs may originate through active secretion or/and passive leakage, as 42.68% of all significant SNV-pQTL proteins identified are actively secreted into the blood at least in one isoform (S18 Table) [67], meaning that more than half of these proteins do not originate from the blood. Similar to our findings, Pietzner et al. [7] recently detected a significant colocalisation of 50.1% of the *cis*-pQTLs with corresponding gene eQTLs using GTEx.

We sought to systematically identify links between proteins and phenotypes by conducting a PheWAS followed by a colocalisation analysis, in order to find signals likely driven by the same causal variant. We then applied MR to significant colocalisation events to assess causality, a strategy recommended by Zuber et al. [17]. As they have highlighted, a positive colocalisation finding typically implies a non-zero MR estimate, the reverse is not generally true [17]. For example, FGF5 plays essential roles in the regulation of cell proliferation, including in cardiac myocytes, and cell differentiation [68]; it has also been associated with cardiac angiogenesis [69]. The *FGF5* locus has been linked to cardiovascular conditions in previous GWASs [62,70]. We detected a *cis* signal for the FGF5 level and associated variants in the region, which overlapped with previous GWAS findings for cardiovascular diseases and medications used to treat them. Our colocalisation and MR results suggest that the FGF5 level shares common causal SNPs with various heart-related conditions and treatments, prioritising it as an interesting target for future follow-up studies. However, the translation of PheWAS results to a molecular level is complicated by the nature of associated disease phenotype. Plasma proteins are potentially more relevant for circulatory diseases where the blood is in contact with the affected tissue, such as in the FGF5 example, rather than for conditions with a limited number of affected tissues.

The availability of the high-quality WGS data also gave us a unique opportunity to investigate the effect of CNVs on protein expression. To the best of our knowledge, one study has previously studied CNVs in this context, focusing only on deletions [15]. We conducted the first comprehensive CNV-based pQTL mapping and identified 12 associations (7 *cis* and 5 *trans*) between plasma proteins and CNVs, including those with a *trans*-association hub CNV in the 3q12.1 region. We further interpreted the CNV-pQTLs using a CNV-tagging SNP approach with external data on a broader range of proteins. This strategy yielded additional CNV-based pQTLs for 79 proteins and determined that the 3q12.1-region hub CNV was associated with 25 proteins. Signals from the SNV and CNV analyses overlapped for nine proteins, which constitute interesting loci where QTL associations were likely driven by CNVs, rather than SNVs. This emphasises the value of the CNV data, especially if the purpose is to prioritise causal genetic variation underlying the pQTL signal. None of the associations reported by Png et al. [15] were replicated in this study, possibly because there was only a partial overlap between the assayed protein sets, differences between cohorts (European ancestry vs a Greek population isolate with population-specific CNVs) [71], and differences in the approach used for CNV detection.

As an example, we outline IL-18, a pro-inflammatory cytokine that plays important roles in natural killer cell activation and the T-helper 1 response [72]. We found that a CNV on chromosome 5 overlapping with *NAIP* has *trans* effects on the IL18 protein level and a *cis* effect on the *NAIP* gene expression level, but there is no significant effect on the *IL18* gene expression. The *NAIP* eQTL signal was stronger than the IL18 pQTL signal, suggesting that the CNV affects the protein level through gene expression. As the NAIP level was not measured in our cohort, it remains unclear whether the main effect of the CNV is on NAIP. To our knowledge, there are no previous studies analysing the effect of genetic variants on NAIP level. NAIP is an anti-apoptotic protein and sensor component of the NLRC4 inflammasome that protects against bacterial pathogens, and NAIP-NLRC4 inflammasome activation has been reported to lead to elevated IL-18 expression in enterocytes and monocyte-derived macrophages [73]. This example highlights the importance of including structural variants in addition to SNVs in studies of the genetic basis of molecular traits, as also exemplified by the CNV-tagging SNP approach.

We identified 19 significant rare variant effects on the levels of seven proteins that would not have been detected by the SNV pQTL analysis alone. Gene-based pQTL analyses of rare variants constitute an emerging approach [10–13], and no golden standard for their performance has been established, making the replication of findings difficult. Previous studies indicate that few proteins are driven by rare variants [11–13]. Kierczak et al. [13] detected *cis*-region rare variant associations for four proteins (CTSZ, CYR61, GDF-15 and PON3) and *trans* associations of rare *GAL3ST2* variants affecting the MUC16 level, the effect also detected in our study; they used a maximum MAF threshold 0.0239, whereas we used a standard conservative threshold of 0.01. The significant rare variant associations detected in our study were not reported in the largest gene-based rare variant pQTL study conducted to date which included three isolated European cohorts with a total sample size of *n*=4,422 [12]. As an example, we found a rare-variant effect on GDF-15, which regulates food intake, energy expenditure and body weight in response to metabolic and toxin-induced stress [74–76]. The most significant association with the GDF-15 level was a *trans* association with rare variants in *JAKMIP1*, associated with type 2 diabetes and medications used to treat it [77–79]. Additionally, GDF-15 has been reported to be involved in inflammation, metabolism and cancer [80], and recent findings support its role as a biomarker of metabolic stress [81]. Whereas we detected rare variant *trans* associations emanating from GDF-15 for nine proteins, only SNP-based *cis* associations with GDF-15 itself have been identified in previous pQTL studies [9,81]. This demonstrates that gene-based rare variant pQTLs complement single variant analyses and help to unravel novel biologically interpretable associations.

Our study has several limitations. First, the sample size was small relative to those of recent pQTL studies, which made the detection of *trans* effects with greater multiple-testing burden and weak effects of common and rare variants more difficult. Rare genetic variants tend to have greater population specificity [82], making replication of findings from rare variant analyses more difficult. Same applies to common CNVs we reported in our pQTL analyses; structural variants are currently understudied in terms of pQTL detection, limiting replicability. Second, most pQTL studies have been conducted using serum or plasma measurements from blood samples [4,6,8,39] and only a limited number of studies has involved the examination of liver and brain tissue–specific pQTLs [83,84]. Therefore, it is often challenging to understand whether observed pQTL effects manifest in the blood cells or reflect the regulation happening in some distal tissue. Finally, although we showed that CNVs affect plasma protein levels, to our knowledge no large-scale CNV-based association database is currently available to overlap the identified CNV-pQTL associations with CNV-phenotype associations. However, CNV-tagging SNPs could be used as a proxy method to assess the effect of CNVs on complex traits.

In conclusion, we have generated a comprehensive pQTL resource and interpreted it by using eQTL, as well as publicly available GWAS data. We have demonstrated the importance of including structural variants in addition to SNVs, to fully characterise the genetic background of plasma proteins and their links to health-related phenotypes.

## Acknowledgements

The authors would like to thank Hanna Maria Kariis for helpful comments on the manuscript. Members of the Estonian Biobank Research Team include Andres Metspalu, Lili Milani, Tõnu Esko, Reedik Mägi, Mari Nelis and Georgi Hudjašov. Data analyses with Estonian datasets were carried out in part in the High-Performance Computing Center of University of Tartu.

## Supporting information

**S1 Table. Full list of the Olink proteins in the study.** Columns are ‘OLINK’: the protein name based on Olink internal naming scheme; ‘PANEL’: Olink panel for the protein; ‘UNIPROT_Olink’: UniProtID of the protein; ‘HGNC’: HUGO gene naming consortium symbol for the protein; ‘LOD_QC’: limit of detection (LOD) quality control assessment. LOD was used as a quality control step, each protein with samples >20% LOD was flagged as fail; ‘Alternative_UNIPROT’: alternative UniProtID of the protein if available.

**S2 Table. List of pQTLs (linkage disequilibrium clumped).** List of lead variants for each protein following linkage disequilibrium (LD) clumping, together with replication information. Variants within a 1 Mb window of the lead pQTL with the LD thresholds of R*^2^* = 0.1 and *P* < 5 × 10^−8^ were clumped together. Whole-genome sequenced genotypes of the pQTL cohort were used as LD reference. Columns are ‘gene’: HUGO gene naming consortium symbol for the protein; ‘Uniprot’: UniProtID of the protein; ‘panel’: Olink panel for the protein; ‘chr_pos’: genomic coordinates for the pQTL variant (hg19); ‘locus’: pQTL association locus; ‘variant’: variant name in the format of genomic coordinates (hg19) and alphabetically ordered alleles; ‘rsid’: rsID (if missing, then genomic coordinates in hg19); ‘A1’: the reference allele in the Estonian Biobank; ‘A2’: the effect allele in the Estonian Biobank; ‘MAF’: minor allele frequency; ‘p-value’: pQTL association *p*-value; ‘beta’: the pQTL effect size; ‘SE’: the standard error of the pQTL effect size; ‘type’: pQTL association signal type, associations within 1Mb upstream or downstream of the transcription start site (TSS) of the corresponding protein-coding genes are *cis* and further away *trans*; ‘distance’: the distance from the TSS for *cis* associations in bp; ‘effect’: the functional annotation of a pQTL; ‘LD R^2^>0.8 PAV variant’: protein-altering variants in linkage disequilibrium (R^2^>0.8) with detected pQTL. For replication studies prefix in column names indicates the name of the study, referring to Pietzner et al. [7], Sun et al. [4], Suhre et al. [39] and Folkersen et al. [38]. Columns are ‘Pietzner_replication’: the pQTL replication in the TRUE (replicating) and FALSE (not replicating) manner; ‘Pietzner_EA’: the effect allele of pQTL in the Pietzner et al.; ‘Pietzner_OA’: the other allele in the Pietzner et al.; ‘Pietzner_effect’: the effect size of pQTL in the Pietzner et al.; ‘Pietzner_se’: the standard error of effect size in the Pietzner et al.; ‘Pietzner_pval’: the *p*-value of pQTL association in the Pietzner et al.; ‘Pietzner_qval_FDR’: the Benjamini-Hochberg FDR-corrected *q*-value of the Pietzner et al. pQTL analysis; ‘Pietzner_n’: the sample size in the Pietzner et al. for the variant; ‘Sun_replication’: the pQTL replication in the TRUE (replicating) and FALSE (not replicating) manner; ‘Sun_EA’: the effect allele of pQTL in the Sun et al; ‘Sun_effect’: the effect size of pQTL in the Sun et al.; ‘Sun_se’: the standard error of effect size in the Sun et al.; ‘Sun_pval’: the *p*-value of pQTL association in the Sun et al.; ‘Sun_qval_FDR’: the Benjamini-Hochberg FDR-corrected *q*-value of the Sun et al. pQTL analysis; ‘Sun_n’: the sample size in the Sun et al. for the variant; ‘Suhre_replication’: the pQTL replication in the TRUE (replicating) and FALSE (not replicating) manner; ‘Suhre_EA’: the effect allele of pQTL in the Suhre et al.; ‘Suhre_effect’: the effect size of pQTL in the Suhre et al.; ‘Suhre_se’: the standard error of effect size in the Suhre et al.; ‘Suhre_pval’: the *p*-value of pQTL association in the Suhre et al.; ‘Suhre_qval_FDR’: the Benjamini-Hochberg FDR-corrected *q*-value of the Suhre et al. pQTL analysis; ‘Suhre_n’: the sample size in the Suhre et al. for the variant; ‘Folkersen_replication’: the pQTL replication in the TRUE (replicating) and FALSE (not replicating) manner; ‘Folkersen_EA’: the effect allele of pQTL in the Folkersen et al.; ‘Folkersen_effect’: the effect size of pQTL in the Folkersen et al.; ‘Folkersen_se’: the standard error of effect size in the Folkersen et al.; ‘Folkersen_pval’: the *p*-value of pQTL association in the Folkersen et al.; ‘Folkersen_qval_FDR’: the Benjamini-Hochberg FDR-corrected *q*-value of the Folkersen et al. pQTL analysis; ‘Folkersen_n’: the sample size in the Folkersen et al. for the variant. In case of Pietzner et al., alleles for indels are referred as ‘D’ for deletion and ‘I’ for insertion.

**S3 Table. List of the eQTL Catalogue resources.** Columns are ‘Study’: the consortium or the publication for the dataset; ‘Publication’: the citation of dataset publication; ‘Funding’: the funding for generating the dataset.

**S4 Table. List of studied complex traits extracted from the Medical Research Council (MRC) Integrative Epidemiology Unit (IEU) OpenGWAS database.** Columns are ‘ID’: internal naming identification for a complex trait GWAS in the MRC IEU OpenGWAS database; ‘Trait’: the full name of the complex trait; ‘n_cases/n_controls’: number of cases/number of controls for the study; ‘Publication/Author’: the consortium or the publication that generated complex trait GWAS results; ‘Funding/Acknowledgements’: funding and acknowledgements marked by the consortium or by the publication.

**S5 Table. List of the corresponding eQTLs.** Columns are ‘variant’: rsID (if missing, then genomic coordinates in hg19); ‘protein’: HUGO gene naming consortium symbol for the protein; ‘Uniprot’: UniProtID of the protein; ‘panel’: Olink panel for the protein; ‘pQTL_pval’: pQTL association *p*-value; ‘pQTL_beta’: pQTL effect size; ‘pQTL_se’: the standard error of the pQTL effect size; ‘type’: pQTL association signal type, associations within 1Mb upstream or downstream of the transcription start site (TSS) of the corresponding protein-coding genes are *cis* and further away *trans*; ‘gene’: HUGO gene naming consortium symbol for the protein for tested gene; Ensembl’: Ensembl (GRCh37) gene ID for tested gene; ‘eQTL_pval’: eQTL association *p*-value; ‘eQTL_beta’: eQTL effect size; ‘eQTL_se’: the standard error of the eQTL effect size; ‘eQTL_qFDR’: the Benjamini-Hochberg FDR-corrected *q*-value of the eQTL analysis.

**S6 Table. (A) Results of the fine-mapping analysis and (B) an overview of variants within each credible set having the highest posterior inclusion probability (PIP).** (A) Columns are ‘trait’: HUGO gene naming consortium symbol for the protein; ‘panel’: Olink panel for the protein; ‘region’: genetic coordinates for the fine-mapping region (GRCh37); ‘locus’: locus and loci in the fine-mapping analysis (GRCh37); ‘credible set’: the number of identified credible sets; ‘size’: the number of genetic variants belonging to the specific credible set; ‘type’: the pQTL association signal type in the primary pQTL analysis (S2 Table). (B) Columns are ‘trait’: HUGO gene naming consortium symbol for the protein; ‘chromosome’: the chromosome of the fine-mapped variant (GRCh37), ‘credible set’: the number of the identified credible set; ‘Fine-mapped variant (GRCh37)’: variant in the format chromosome: region and alleles ordered alphabetically (GRCh37); ‘PIP’: the posterior inclusion probability of the variant; ‘association p-value’: the p-value of the variant in the pQTL analysis; ‘LD (r²) with sentinel SNP’: the linkage disequilibrium of the fine-mapped variant with the primary pQTL identified in the pQTL analysis; ‘Distance (kb) with sentinel SNP (GRCh37)’: genetic distance in kb between fine-mapped variant and pQTL identified in the region in the primary analysis.

**S7 Table. Regulatory information for the pQTLs extracted from the RegulomeDB.** RegulomeBD classifies SNPs into classes based on the combinatorial presence/absence status of functional categories, including transcription factors binding sites, DNAase hypersensitivity regions, and promoter regions. Columns are ‘chrom’: the chromosome of the pQTL variant (hg19); ‘start’: start coordinates of the queried variant (hg19); ‘end’: end coordinates of the queried variant (hg19): ‘rsids’: rsID for the queried variant; ‘probability’: probability score ranging from 0 to 1, with 1 being the most likely regulatory variant; ‘ranking’: ranking based on RegulomeDB internal scoring scheme that takes into account supporting data. Categories included in the table are ‘1d’: eQTL + TF binding + any motif + DNase peak; ‘1f’: ‘eQTL + TF binding / DNase peak’; ‘2a’: TF binding + matched TF motif + matched DNase Footprint + DNase peak; ‘2b’: TF binding + any motif + DNase Footprint + DNase peak; ‘2c’: TF binding + matched TF motif + DNase peak; ‘3a’: TF binding + any motif + DNase peak; ‘4’: TF binding + DNase peak; ‘5’: TF binding or DNase peak; ‘6’: Motif hit; ‘7’: Other.

**S8 Table. List of colocalising pQTL**–**eQTL events.** Columns are ‘pQTL_lead_SNP_HG19’: genomic coordinates for the primary pQTL (hg19); ‘pQTL_lead_SNP_HG38’: genomic coordinates for the primary pQTL (hg38); ‘pQTL_Uniprot’: UniProtID for the protein; ‘pQTL_Gene_Ensembl’: Ensembl gene ID; ‘pQTL_Gene_Name’: HUGO gene naming consortium symbol for the protein; ‘pQTL_Gene_Loc_HG38’: pQTL gene genomic coordinates (hg38); ‘pQTL_Cis_Trans’: the association type for the pQTL in the primary analysis, either local *cis* or distal *trans*; ‘eQTL_Gene_Ensembl’: Ensembl gene ID for the tested eQTL gene; ‘eQTL_Gene_Name’: HUGO gene naming consortium symbol for the eQTL gene; ‘eQTL_Gene_Loc_HG38’: eQTL gene genomic coordinates (hg38); ‘eQTL_Trait’: ID of the molecular trait used for QTL mapping, depending on the quantification method used, this can be either a gene id, exon id, transcript id or a txrevise promoter, splicing or 3’end event id; ‘eQTL_Dataset’: eQTL dataset name and tested and tissue or cell type and trait quantification; ‘Study’: the study or the consortium of the eQTL data; ‘eQTL_Data_Type’: quantification type in the eQTL Catalogue as either gene expression, exon expression, transcript usage or txrevise event usage; ‘Tissue_Cells’: tissue or cell type for the eQTL; ‘nsnps’: the number of SNPs included in the genetic region of the colocalisation analysis; ‘PP.H0.abf’: posterior probability of no association with either trait; ‘PP.H1.abf’: posterior probability of association with pQTL but not eQTL; ‘PP.H2.abf’: posterior probability of association with eQTL but not pQTL; ‘PP.H3.abf’: posterior probability of association with both traits but at separate causal variants; ‘PP.H4.abf’: posterior probability of association with both traits at a shared causal variant.

**S9 Table. List of pQTLs from the metabolites PheWAS.** Columns are ‘snp’: the queried SNP rsID; ‘rsid’: the queried SNP rsID; ‘hg19_coordinates’: genomic coordinates for the queried SNP (hg19); ‘hg38_coordinates’: genomic coordinates for the queried SNP (hg38); ‘a1’: the effect allele for the queried SNP; ‘a2’: the non-effect allele for the queried SNP; ‘trait’: the metabolite phenotype; ‘efo’: corresponding EFO ontology term for the metabolite phenotype; ‘study’: the name of the consortium or lead author of the study; ‘pmid’: the PubMed ID; ‘ancestry’: the ancestry of the study; ‘year’: the year the study was published; ‘beta’: the association between the trait and the SNP expressed per additional copy of the effect allele (odds ratios are given on the log-scale); ‘se’: the standard error of beta; ‘p’: the *p*-value; ‘direction’: the direction of association with respect to the effect allele; ‘n’: the number of individuals; ‘n_studies’: the number of studies; ‘unit’: the unit of analysis (IVNT stands for inverse normally rank transformed phenotype); ‘dataset’: the dataset ID as the first author or the consortium.

**S10 Table. List of pQTLs from the epigenetics PheWAS.** Columns are ‘snp’: the queried SNP rsID; ‘rsid’: the queried SNP rsID; ‘hg19_coordinates’: genomic coordinates for the queried SNP (hg19); ‘hg38_coordinates’: genomic coordinates for the queried SNP (hg38); ‘a1’: the effect allele for the queried SNP; ‘a2’: the non-effect allele for the queried SNP; ‘trait’: the epigenetics phenotype; ‘efo’: corresponding EFO ontology term for the epigenetics phenotype; ‘study’: the name of the consortium or lead author of the study; ‘pmid’: the PubMed ID; ‘ancestry’: the ancestry of the study; ‘year’: the year the study was published; ‘tissue’: the tissue in which the gene expression was measured; ‘marker’: the epigenetic marker measured; ‘location’ the location of epigenetic marker (hg19); ‘beta’: the association between the trait and the SNP expressed per additional copy of the effect allele (odds ratios are given on the log-scale); ‘se’: the standard error of beta; ‘p’: the *p*-value; ‘direction’: the direction of association with respect to the effect allele; ‘n’: the number of individuals; ‘n_studies’: the number of studies; ‘unit’: the unit of analysis (IVNT stands for inverse normally rank transformed phenotype); ‘dataset’: the dataset ID as the first author or the consortium.

**S11 Table. List of pQTLs from the PheWAS.** Columns are ‘snp’: the queried SNP rsID; ‘rsid’: the queried SNP rsID; ‘ref_hg19_coordinates’: the queried SNP genomic coordinates (hg19); ‘ref_hg38_coordinates’: the queried SNP genomic coordinates (hg38); ‘ref_a1’: the effect allele for the queried SNP; ‘ref_a2’: the non-effect allele for the queried SNP; ‘rsid’: the rsID for the proxy SNP; ‘hg19_coordinates’: genomic coordinates for the proxy SNP (hg19); ‘hg38_coordinates’; genomic coordinates for the proxy SNP (hg38); ‘rsid’: the rsID for the proxy SNP; ‘ref_a1’: the effect allele for the proxy SNP; ‘ref_a2’: the non-effect allele for the proxy SNP; ‘proxy’: an indicator variable which equals 0 if the proxy SNP is the input SNP and 1 otherwise; ‘r2’: the *r*^2^ between the input SNP and the proxy SNP based on the phased haplotypes from 1000 Genomes; ‘dprime’: the D’ between the input SNP and the proxy SNP based on the phased haplotypes from 1000 Genomes; ‘trait’: the phenotype; ‘efo’: corresponding EFO ontology term for the phenotype; ‘study’: the name of the consortium or lead author of the study; ‘pmid’: the PubMed ID; ‘ancestry’: the ancestry of the study; ‘year’: the year the study was published; ‘beta’: the association between the trait and the SNP expressed per additional copy of the effect allele (odds ratios are given on the log-scale); ‘se’: the standard error of beta; ‘p’: the *p*-value; ‘direction’: the direction of association with respect to the effect allele; ‘n’: the number of individuals; ‘n_cases’: the number of cases; ‘n_controls’: the number of controls; ‘n_studies’: the number of studies; ‘unit’: the unit of analysis (IVNT stands for inverse normally rank transformed phenotype); ‘dataset’: the dataset ID as the first author or the consortium.

**S12 Table. Results from the pQTL**–**complex trait colocalisation analysis.** Columns are ‘Protein’: HUGO gene naming consortium symbol for the protein, ‘ID’: internal identification for complex trait used in the MRC CEU OpenGWAS database; ‘Study’: the name of the consortium/biobank or the first author of the study; ‘Trait’: the full naming of the complex trait in the MRC CEU OpenGWAS database; ‘nsnps’: the number of SNPs included in the genetic region of the colocalisation analysis; ‘PP.H0.abf’: posterior probability of no association with either trait (if PP_0_ > 0.8); ‘PP.H1.abf’: posterior probability of association with pQTL but not complex trait (if PP_1_ > 0.8); ‘PP.H2.abf’: posterior probability of association with complex trait but not pQTL (PP_2_ > 0.8); ‘PP.H3.abf’: posterior probability of association with both traits but at separate causal variants (if PP_3_ > 0.8); ‘PP.H4.abf’: posterior probability of association with both traits at a shared causal variant (if PP_4_ > 0.8).

**S13 Table. Results from the pQTL–complex trait Mendelian randomisation analysis.** Columns are ‘Protein’: HUGO gene naming consortium symbol for the protein used as exposure; ‘Trait’: the complex trait used as an outcome; ‘Full trait name’: the full naming of the complex trait in the MRC CEU OpenGWAS database; ‘ID’: internal identification for complex trait used in the MRC CEU OpenGWAS database; ‘Study’: the name of the consortium/biobank or the first author of the study; ‘Test’: the method used to conduct Mendelian randomisation (MR), for single variant based exposure traits Wald test and for multiple variants based exposure traits inverse variance weighted (IVW) regression; ‘nSNP’: the number of genetic variants used as instrumental variables (IV) in exposure traits for the MR analysis; ‘b’: the causal effect estimate of the protein (exposure) on the complex trait (outcome); ‘se’: the standard error of the causal effect estimate; ‘pval’: the *p*-value of the MR analysis; ‘qFDR’: the Benjamini-Hochberg FDR-corrected *q*-value of the MR analysis.

**S14 Table. List of significant (*P* < 1.48 × 10^−8^) associations from the rare variant gene-based pQTL analysis.** Columns are ‘Uniprot’: UniProtID of the protein; ‘Protein’: HUGO gene naming consortium symbol for the protein; ‘chr’: the chromosome of the associated gene (GRCh37), ‘beg’: the start coordinates of the gene (GRCh37); ‘end’: the end coordinates of the gene (GRCh37); ‘marker_id’: the genetic location of the associated gene, including chromosome, start and end coordinates, and HGNC gene symbol for it (GRCh37); ‘NS’: the number of phenotyped samples with non-missing genotypes; ‘FRAC_WITH_RARE’: the fraction of individuals carrying rare variants below the maximum of minor allele frequency threshold (MAF < 0.01); ‘NUM_ALL_VARS’: the number of all variants defining the group, meaning all genetic variants located within the tested gene; ‘NUM_PASS_VARS’: the number of variants passing the minimum of MAF (0.0000001), the minimum of minor allele count (1), the maximum of MAF (0.01) and minimum of call rate (0.5) thresholds; ‘NUM_SING_VARS’: the number of singletons among variants in ‘NUM_PASS_VARS’; ‘PVALUE’: the *p*-value of the burden test; ‘QSTAT’: the *Q* statistic of the burden test; ‘TYPE’: the association type, *cis* is if the association is with the protein-encoding gene itself and otherwise *trans*; ‘eQTL_NS’: the number of phenotyped samples with non-missing genotypes for gene expression; ‘eQTL_FRAC_WITH_RARE’: the fraction of individuals carrying rare variants below the maximum of minor allele frequency threshold (MAF < 0.01) for gene expression; ‘eQTL_NUM_ALL_VARS’: the number of all variants defining the group, all genetic variants located within the tested gene for gene expression; ‘eQTL_NUM_PASS_VARS’: the number of variants passing the minimum of MAF (0.0000001), the minimum of minor allele count (1), the maximum of MAF (0.01) and minimum of call rate (0.5) thresholds for gene expression; ‘eQTL_NUM_SING_VARS’: the number of singletons among variants in ‘eQTL_NUM_PASS_VARS’ for gene expression; ‘eQTL_PVALUE’: the *p*-value of the burden test for gene expression; ‘eQTL_QSTAT’: the *Q* statistic of the burden test for gene expression; ‘eQTL_QVALUE_FDR’: the Benjamini-Hochberg FDR-corrected *q*-value of the eQTL analysis.

**S15 Table. (A) List of significant CNV-pQTLs and (B) CNV pQTL-eQTL Spearman correlations and MR results.** (A) Columns are ‘CNV’: the genetic location of the CNV in the format chromosome:start-end (GRCh37); ‘Chr’: the chromosome CNV is located on (GRCh37); ‘Start’: the start coordinates of the CNV (GRCh37); ‘End’: the end coordinates of the CNV (GRCh37); ‘Uniprot’: UniProtID of the protein; ‘Array’: Olink panel for the protein; ‘Gene’: HUGO gene naming consortium symbol for the protein; ‘Type’: the pQTL association type, if the CNV association is in the proximity of the protein-encoding gene, the association is *cis* and otherwise *trans*; ‘P-value (pQTL)’: the CNV pQTL association *p*-value; ‘P-value (eQTL same gene)’: the *p*-value from the CNV eQTL analysis for the pQTL gene (for heterodimer the specific subunit is in the brackets); ‘P-value (eQTL other gene)’: the *p*-value from the CNV eQTL analysis for not pQTL gene and in the brackets in the associated gene; ‘CNV overlap with a gene’: CNV overlap with a gene and gene symbol is in the brackets, for heterodimer, overlap with subunit is marked; ‘Number of copies’: the possible number of alleles detected for the CNV in the Estonian population; ‘Allele frequency’: the frequency of the CNV based on the number of copies corresponding in the column ‘Number of copies’. (B) Columns are ‘CNV’: the genetic location of the CNV in the format chromosome:start-end (GRCh37); ‘gene (RNAseq)’: HUGO gene naming consortium symbol for the gene; ‘protein (Olink)’: HUGO gene naming consortium symbol for the protein; ‘R (Spearman)’: Spearman’s rank correlation coefficient for gene expression versus protein expression; ‘Z (MR)’: Z score as causal effect estimate from the CNV eQTL and CNV pQTL MR analysis; ‘P (MR)’: *p*-value from the CNV eQTL and CNV pQTL MR analysis. *Reference for these values is a whole-genome sequenced cohort of 2,273 individuals in the Estonian Biobank.

**S16 Table. List of significant CNV-eQTLs.** Columns are ‘CNV’: genetic coordinates of the tested CNV in the format of chromosome:start-end (GRCh37); ‘gene’: HUGO gene naming consortium symbol for the trait; ‘ensembl’: Ensembl transcript ID for the tested trait; ‘beta’: estimate of the effect size; ‘t-stat’: t-statistic of the association; ‘p-value’: *p*-value of the association; ‘FDR’: Benjamini-Hochberg procedure corrected association *q*-value.

**S17 Table. List of pQTLs identified in the SNP-tagged CNV analysis.** Single variant pQTL results originate from the Sun et al. 2022 study [9]. Columns are ‘chr’: the chromosome (hg19); ‘position’: the position of the SNP (hg19); ‘rsID’: the rsID of the pQTL SNP; ‘A1’: the reference allele; ‘A2’: the tested allele; ‘target’: HUGO gene naming consortium symbol for the protein; ‘cis_trans’: the association type in the original Sun et al. [9] pQTL mapping (either *cis* or *trans*); ‘A2_freq_discovery’: the frequency of the tested SNP in the Sun et al. [9] discovery cohort; ‘A2_freq_replication’: the frequency of the tested SNP in the Sun et al. [9] replication cohort; ‘A2_freq_Est’: the frequency of the tested SNP in the Estonian Biobank; ‘maxR2’: the R^2^ of the linkage disequilibrium between Sun et al. [9] pQTL SNP and the Estonian Biobank CNV; ‘maxR2_CNV’: the CNV tagged by SNPs coordinates (hg19); ‘frequency (deletion/duplication)’: the frequency of the CNV in the Estonian Biobank; ‘maxR2_CNV_Impact’: the classification of the most likely impact of the SNP tagging the CNV; ‘maxR2_CNV_Consequence’: the most likely consequence of the SNP tagging the CNV.

**S18 Table. List of secretion locations for the proteins with significant results from the pQTL analysis.** Columns are ‘Protein’: the HGNC gene symbol for the protein; ‘location’: the location of proteins; ‘CNV’: CNV pQTL association detection; ‘rare’: rare variant gene-based pQTL association detection.

## Notes

### Competing Interest Statement

The authors have declared no competing interest.

## References

1. MacArthur J, Bowler E, Cerezo M, Gil L, Hall P, Hastings E, et al. The new NHGRI-EBI Catalog of published genome-wide association studies (GWAS Catalog). Nucleic Acids Res. 2017 04;45(D1):D896–901.

2. Maurano MT, Humbert R, Rynes E, Thurman RE, Haugen E, Wang H, et al. Systematic Localization of Common Disease-Associated Variation in Regulatory DNA. Science. 2012 Sep 7;337(6099):1190–5.

3. Geyer PE, Holdt LM, Teupser D, Mann M. Revisiting biomarker discovery by plasma proteomics. Mol Syst Biol [Internet]. 2017 Sep 26 [cited 2020 Dec 3];13(9). Available from: https://www.ncbi.nlm.nih.gov/pmc/articles/PMC5615924/

4. Sun BB, Maranville JC, Peters JE, Stacey D, Staley JR, Blackshaw J, et al. Genomic atlas of the human plasma proteome. Nature. 2018 Jun;558(7708):73–9.

5. Emilsson V, Ilkov M, Lamb JR, Finkel N, Gudmundsson EF, Pitts R, et al. Co-regulatory networks of human serum proteins link genetics to disease. Science. 2018 Aug 24;361(6404):769–73.

6. Folkersen L, Gustafsson S, Wang Q, Hansen DH, Hedman ÅK, Schork A, et al. Genomic and drug target evaluation of 90 cardiovascular proteins in 30,931 individuals. Nat Metab. 2020 Oct;2(10):1135–48.

7. Pietzner M, Wheeler E, Carrasco-Zanini J, Cortes A, Koprulu M, Wörheide MA, et al. Mapping the proteo-genomic convergence of human diseases. Science. 2021 Nov 12;374(6569):eabj1541.

8. Ferkingstad E, Sulem P, Atlason BA, Sveinbjornsson G, Magnusson MI, Styrmisdottir EL, et al. Large-scale integration of the plasma proteome with genetics and disease. Nat Genet. 2021 Dec 2;1–10.

9. Sun BB, Chiou J, Traylor M, Benner C, Hsu YH, Richardson TG, et al. Genetic regulation of the human plasma proteome in 54,306 UK Biobank participants [Internet]. bioRxiv; 2022 [cited 2022 Jun 20]. p. 2022.06.17.496443. Available from: https://www.biorxiv.org/content/10.1101/2022.06.17.496443v1

10. Solomon T, Lapek JD, Jensen SB, Greenwald WW, Hindberg K, Matsui H, et al. Identification of Common and Rare Genetic Variation Associated With Plasma Protein Levels Using Whole-Exome Sequencing and Mass Spectrometry. Circ Genomic Precis Med. 2018;11(12):e002170.

11. Gilly A, Park YC, Png G, Barysenka A, Fischer I, Bjørnland T, et al. Whole-genome sequencing analysis of the cardiometabolic proteome. Nat Commun. 2020 Dec 10;11(1):6336.

12. Gilly A, Klaric L, Park YC, Png G, Barysenka A, Marsh JA, et al. Gene-based whole genome sequencing meta-analysis of 250 circulating proteins in three isolated European populations. Mol Metab. 2022 Apr 30;101509.

13. Kierczak M, Rafati N, Höglund J, Gourlé H, Lo Faro V, Schmitz D, et al. Contribution of rare whole-genome sequencing variants to plasma protein levels and the missing heritability. Nat Commun. 2022 May 9;13(1):2532.

14. Dhindsa RS, Burren OS, Sun BB, Prins BP, Matelska D, Wheeler E, et al. Influences of rare protein-coding genetic variants on the human plasma proteome in 50,829 UK Biobank participants [Internet]. bioRxiv; 2022 [cited 2022 Oct 13]. p. 2022.10.09.511476. Available from: https://www.biorxiv.org/content/10.1101/2022.10.09.511476v1

15. Png G, Suveges D, Park YC, Walter K, Kundu K, Ntalla I, et al. Population-wide copy number variation calling using variant call format files from 6,898 individuals. Genet Epidemiol. 2020;44(1):79–89.

16. Zheng J, Haberland V, Baird D, Walker V, Haycock PC, Hurle MR, et al. Phenome-wide Mendelian randomization mapping the influence of the plasma proteome on complex diseases. Nat Genet [Internet]. 2020 Sep 7 [cited 2020 Sep 8]; Available from: http://www.nature.com/articles/s41588-020-0682-6

17. Zuber V, Grinberg NF, Gill D, Manipur I, Slob EAW, Patel A, et al. Combining evidence from Mendelian randomization and colocalization: Review and comparison of approaches. Am J Hum Genet [Internet]. 2022 Apr 21 [cited 2022 May 2];0(0). Available from: https://www.cell.com/ajhg/abstract/S0002-9297(22)00149-5

18. Kerimov N, Hayhurst JD, Peikova K, Manning JR, Walter P, Kolberg L, et al. A compendium of uniformly processed human gene expression and splicing quantitative trait loci. Nat Genet. 2021 Sep 6;1–10.

19. Leitsalu L, Haller T, Esko T, Tammesoo ML, Alavere H, Snieder H, et al. Cohort Profile: Estonian Biobank of the Estonian Genome Center, University of Tartu. Int J Epidemiol. 2015 Aug;44(4):1137–47.

20. Mitt M, Kals M, Pärn K, Gabriel SB, Lander ES, Palotie A, et al. Improved imputation accuracy of rare and low-frequency variants using population-specific high-coverage WGS-based imputation reference panel. Eur J Hum Genet EJHG. 2017;25(7):869–76.

21. Handsaker RE, Van Doren V, Berman JR, Genovese G, Kashin S, Boettger LM, et al. Large multiallelic copy number variations in humans. Nat Genet. 2015 Mar;47(3):296– 303.

22. Lepamets M, Auwerx C, Nõukas M, Claringbould A, Porcu E, Kals M, et al. Omics-informed CNV calls reduce false positive rate and improve power for CNV-trait associations [Internet]. bioRxiv; 2022 [cited 2022 Jun 9]. p. 2022.02.07.479374. Available from: https://www.biorxiv.org/content/10.1101/2022.02.07.479374v1

23. Assarsson E, Lundberg M, Holmquist G, Björkesten J, Thorsen SB, Ekman D, et al. Homogenous 96-Plex PEA Immunoassay Exhibiting High Sensitivity, Specificity, and Excellent Scalability. PLOS ONE. 2014 Apr 22;9(4):e95192.

24. Bolger AM, Lohse M, Usadel B. Trimmomatic: a flexible trimmer for Illumina sequence data. Bioinforma Oxf Engl. 2014 Aug 1;30(15):2114–20.

25. Andrews S. FastQC: Aquality control tool for high throughput sequence data 2010. Available from: [Internet]. [cited 2023 Mar 7]. Available from: https://www.bioinformatics.babraham.ac.uk/projects/fastqc/

26. Dobin A, Davis CA, Schlesinger F, Drenkow J, Zaleski C, Jha S, et al. STAR: ultrafast universal RNA-seq aligner. Bioinforma Oxf Engl. 2013 Jan 1;29(1):15–21.

27. Robinson MD, Oshlack A. A scaling normalization method for differential expression analysis of RNA-seq data. Genome Biol. 2010;11(3):R25.

28. Robinson MD, McCarthy DJ, Smyth GK. edgeR: a Bioconductor package for differential expression analysis of digital gene expression data. Bioinforma Oxf Engl. 2010 Jan 1;26(1):139–40.

29. Lepik K, Annilo T, Kukuškina V, Consortium eQTLGen, Kisand K, Kutalik Z, et al. C-reactive protein upregulates the whole blood expression of CD59 - an integrative analysis. PLOS Comput Biol. 2017 Sep 18;13(9):e1005766.

30. Kang HM, Sul JH, Service SK, Zaitlen NA, Kong SY, Freimer NB, et al. Variance component model to account for sample structure in genome-wide association studies. Nat Genet. 2010 Apr;42(4):348–54.

31. Chang CC, Chow CC, Tellier LC, Vattikuti S, Purcell SM, Lee JJ. Second-generation PLINK: rising to the challenge of larger and richer datasets. GigaScience. 2015;4:7.

32. Boyle AP, Hong EL, Hariharan M, Cheng Y, Schaub MA, Kasowski M, et al. Annotation of functional variation in personal genomes using RegulomeDB. Genome Res. 2012 Sep 1;22(9):1790–7.

33. Gao X, Starmer J, Martin ER. A multiple testing correction method for genetic association studies using correlated single nucleotide polymorphisms. Genet Epidemiol. 2008;32(4):361–9.

34. Kettunen J, Demirkan A, Würtz P, Draisma HHM, Haller T, Rawal R, et al. Genome-wide study for circulating metabolites identifies 62 loci and reveals novel systemic effects of LPA. Nat Commun. 2016 Mar 23;7(1):11122.

35. Lê S, Josse J, Husson F. FactoMineR: An R Package for Multivariate Analysis. J Stat Softw. 2008 Mar 18;25(1):1–18.

36. Harrow J, Frankish A, Gonzalez JM, Tapanari E, Diekhans M, Kokocinski F, et al. GENCODE: The reference human genome annotation for The ENCODE Project. Genome Res. 2012 Sep 1;22(9):1760–74.

37. Franz M, Rodriguez H, Lopes C, Zuberi K, Montojo J, Bader GD, et al. GeneMANIA update 2018. Nucleic Acids Res. 2018 02;46(W1):W60–4.

38. Warde-Farley D, Donaldson SL, Comes O, Zuberi K, Badrawi R, Chao P, et al. The GeneMANIA prediction server: biological network integration for gene prioritization and predicting gene function. Nucleic Acids Res. 2010 Jul;38(Web Server issue):W214-220.

39. Wang G, Sarkar A, Carbonetto P, Stephens M. A simple new approach to variable selection in regression, with application to genetic fine mapping. J R Stat Soc Ser B Stat Methodol. 2020;82(5):1273–300.

40. Zou Y, Carbonetto P, Wang G, Stephens M. Fine-mapping from summary data with the “Sum of Single Effects” model. PLOS Genet. 2022 juuli;18(7):e1010299.

41. Di Tommaso P, Chatzou M, Floden EW, Barja PP, Palumbo E, Notredame C. Nextflow enables reproducible computational workflows. Nat Biotechnol. 2017 Apr;35(4):316– 9.

42. Benner C, Havulinna AS, Järvelin MR, Salomaa V, Ripatti S, Pirinen M. Prospects of Fine-Mapping Trait-Associated Genomic Regions by Using Summary Statistics from Genome-wide Association Studies. Am J Hum Genet. 2017 Oct 5;101(4):539–51.

43. Kamat MA, Blackshaw JA, Young R, Surendran P, Burgess S, Danesh J, et al. PhenoScanner V2: an expanded tool for searching human genotype-phenotype associations. Bioinforma Oxf Engl. 2019 01;35(22):4851–3.

44. Staley JR, Blackshaw J, Kamat MA, Ellis S, Surendran P, Sun BB, et al. PhenoScanner: a database of human genotype-phenotype associations. Bioinforma Oxf Engl. 2016 15;32(20):3207–9.

45. Folkersen L, Fauman E, Sabater-Lleal M, Strawbridge RJ, Frånberg M, Sennblad B, et al. Mapping of 79 loci for 83 plasma protein biomarkers in cardiovascular disease. PLoS Genet. 2017 Apr;13(4):e1006706.

46. Suhre K, Arnold M, Bhagwat AM, Cotton RJ, Engelke R, Raffler J, et al. Connecting genetic risk to disease end points through the human blood plasma proteome. Nat Commun. 2017 Feb 27;8(1):14357.

47. Finan Chris, Gaulton Anna, Kruger Felix A., Lumbers R. Thomas, Shah Tina, Engmann Jorgen, et al. The druggable genome and support for target identification and validation in drug development. Sci Transl Med. 2017 Mar 29;9(383):eaag1166.

48. Freshour SL, Kiwala S, Cotto KC, Coffman AC, McMichael JF, Song JJ, et al. Integration of the Drug–Gene Interaction Database (DGIdb 4.0) with open crowdsource efforts. Nucleic Acids Res. 2021 Jan 8;49(D1):D1144–51.

49. Elsworth B, Lyon M, Alexander T, Liu Y, Matthews P, Hallett J, et al. The MRC IEU OpenGWAS data infrastructure [Internet]. 2020 Aug [cited 2022 Jan 21] p. 2020.08.10.244293. Available from: https://www.biorxiv.org/content/10.1101/2020.08.10.244293v1

50. Giambartolomei C, Vukcevic D, Schadt EE, Franke L, Hingorani AD, Wallace C, et al. Bayesian Test for Colocalisation between Pairs of Genetic Association Studies Using Summary Statistics. PLOS Genet. 2014 May 15;10(5):e1004383.

51. Wallace C. Eliciting priors and relaxing the single causal variant assumption in colocalisation analyses. PLOS Genet. 2020 Apr 20;16(4):e1008720.

52. Kasela S, Kisand K, Tserel L, Kaleviste E, Remm A, Fischer K, et al. Pathogenic implications for autoimmune mechanisms derived by comparative eQTL analysis of CD4+ versus CD8+ T cells. PLoS Genet. 2017 Mar;13(3):e1006643.

53. GTEx Consortium. The GTEx Consortium atlas of genetic regulatory effects across human tissues. Science. 2020 Sep 11;369(6509):1318–30.

54. Bretherick AD, Canela-Xandri O, Joshi PK, Clark DW, Rawlik K, Boutin TS, et al. Linking protein to phenotype with Mendelian Randomization detects 38 proteins with causal roles in human diseases and traits. PLOS Genet. 2020 Jun 7;16(7):e1008785.

55. Hemani G, Zheng J, Elsworth B, Wade KH, Haberland V, Baird D, et al. The MR-Base platform supports systematic causal inference across the human phenome. Loos R, editor. eLife. 2018 mai;7:e34408.

56. Hemani G, Tilling K, Smith GD. Orienting the causal relationship between imprecisely measured traits using GWAS summary data. PLOS Genet. 2017 Nov 17;13(11):e1007081.

57. Shabalin AA. Matrix eQTL: ultra fast eQTL analysis via large matrix operations. Bioinformatics. 2012 May 15;28(10):1353–8.

58. Zhu Z, Zhang F, Hu H, Bakshi A, Robinson MR, Powell JE, et al. Integration of summary data from GWAS and eQTL studies predicts complex trait gene targets. Nat Genet. 2016 May 1;48(5):481–7.

59. Hao Z, Lv D, Ge Y, Shi J, Weijers D, Yu G, et al. RIdeogram: drawing SVG graphics to visualize and map genome-wide data on the idiograms. PeerJ Comput Sci. 2020 Jan 20;6:e251.

60. Macdonald-Dunlop E, Klarić L, Folkersen L, Timmers PRHJ, Gustafsson S, Zhao JH, et al. Mapping genetic determinants of 184 circulating proteins in 26,494 individuals to connect proteins and diseases. medRxiv. 2021 Jan 1;2021.08.03.21261494.

61. Astle WJ, Elding H, Jiang T, Allen D, Ruklisa D, Mann AL, et al. The Allelic Landscape of Human Blood Cell Trait Variation and Links to Common Complex Disease. Cell. 2016 Nov;167(5):1415–1429.e19.

62. van der Harst P, Verweij N. Identification of 64 Novel Genetic Loci Provides an Expanded View on the Genetic Architecture of Coronary Artery Disease. Circ Res. 2018 Feb 2;122(3):433–43.

63. Stahl EA, Raychaudhuri S, Remmers EF, Xie G, Eyre S, Thomson BP, et al. Genome-wide association study meta-analysis identifies seven new rheumatoid arthritis risk loci. Nat Genet. 2010 Jun;42(6):508–14.

64. Interleukin-6 Receptor Mendelian Randomisation Analysis (IL6R MR) Consortium, Swerdlow DI, Holmes MV, Kuchenbaecker KB, Engmann JEL, Shah T, et al. The interleukin-6 receptor as a target for prevention of coronary heart disease: a mendelian randomisation analysis. Lancet Lond Engl. 2012 Mar 31;379(9822):1214–24.

65. Suhre K, McCarthy MI, Schwenk JM. Genetics meets proteomics: perspectives for large population-based studies. Nat Rev Genet [Internet]. 2020 Aug 28 [cited 2020 Sep 1]; Available from: http://www.nature.com/articles/s41576-020-0268-2

66. Zhang J, Dutta D, Köttgen A, Tin A, Schlosser P, Grams ME, et al. Plasma proteome analyses in individuals of European and African ancestry identify cis-pQTLs and models for proteome-wide association studies. Nat Genet. 2022 May 2;1–10.

67. Uhlén M, Karlsson MJ, Hober A, Svensson AS, Scheffel J, Kotol D, et al. The human secretome Sci Signal [Internet]. 2019 Nov 26 [cited 2020 Mar 2];12(609). Available from: https://stke.sciencemag.org/content/12/609/eaaz0274

68. Ornitz DM, Xu J, Colvin JS, McEwen DG, MacArthur CA, Coulier F, et al. Receptor specificity of the fibroblast growth factor family. J Biol Chem. 1996 Jun 21;271(25):15292–7.

69. Vatner SF. FGF Induces Hypertrophy and Angiogenesis in Hibernating Myocardium. Circ Res. 2005 Apr 15;96(7):705–7.

70. Nikpay M, Goel A, Won HH, Hall LM, Willenborg C, Kanoni S, et al. A comprehensive 1,000 Genomes-based genome-wide association meta-analysis of coronary artery disease. Nat Genet. 2015 Oct;47(10):1121–30.

71. Panoutsopoulou K, Hatzikotoulas K, Xifara DK, Colonna V, Farmaki AE, Ritchie GRS, et al. Genetic characterization of Greek population isolates reveals strong genetic drift at missense and trait-associated variants. Nat Commun. 2014 Nov 6;5(1):5345.

72. Tominaga K, Yoshimoto T, Torigoe K, Kurimoto M, Matsui K, Hada T, et al. IL-12 synergizes with IL-18 or IL-1beta for IFN-gamma production from human T cells. Int Immunol. 2000 Feb 1;12(2):151–60.

73. Kay C, Wang R, Kirkby M, Man SM. Molecular mechanisms activating the NAIP-NLRC4 inflammasome: Implications in infectious disease, autoinflammation, and cancer. Immunol Rev. 2020;297(1):67–82.

74. Emmerson PJ, Wang F, Du Y, Liu Q, Pickard RT, Gonciarz MD, et al. The metabolic effects of GDF15 are mediated by the orphan receptor GFRAL. Nat Med. 2017 Oct;23(10):1215–9.

75. Hsu JY, Crawley S, Chen M, Ayupova DA, Lindhout DA, Higbee J, et al. Non-homeostatic body weight regulation through a brainstem-restricted receptor for GDF15. Nature. 2017 Oct;550(7675):255–9.

76. Yang L, Chang CC, Sun Z, Madsen D, Zhu H, Padkjær SB, et al. GFRAL is the receptor for GDF15 and is required for the anti-obesity effects of the ligand. Nat Med. 2017 Oct;23(10):1158–66.

77. Mahajan A, Wessel J, Willems SM, Zhao W, Robertson NR, Chu AY, et al. Refining the accuracy of validated target identification through coding variant fine-mapping in type 2 diabetes. Nat Genet. 2018 Apr;50(4):559–71.

78. Wu Y, Byrne EM, Zheng Z, Kemper KE, Yengo L, Mallett AJ, et al. Genome-wide association study of medication-use and associated disease in the UK Biobank. Nat Commun. 2019 Apr;10(1):1891.

79. Vujkovic M, Keaton JM, Lynch JA, Miller DR, Zhou J, Tcheandjieu C, et al. Discovery of 318 new risk loci for type 2 diabetes and related vascular outcomes among 1.4 million participants in a multi-ancestry meta-analysis. Nat Genet. 2020 Jul;52(7):680– 91.

80. Breit SN, Johnen H, Cook AD, Tsai VWW, Mohammad MG, Kuffner T, et al. The TGF-β superfamily cytokine, MIC-1/GDF15: a pleotrophic cytokine with roles in inflammation, cancer and metabolism. Growth Factors Chur Switz. 2011 Oct;29(5):187–95.

81. Lemmelä S, Wigmore EM, Benner C, Havulinna AS, Ong RM, Kempf T, et al. Integrated analyses of growth differentiation factor-15 concentration and cardiometabolic diseases in humans. Janus ED, Barton M, Janus ED, editors. eLife. 2022 Aug 2;11:e76272.

82. Momozawa Y, Mizukami K. Unique roles of rare variants in the genetics of complex diseases in humans. J Hum Genet. 2021 Jan;66(1):11–23.

83. He B, Shi J, Wang X, Jiang H, Zhu HJ. Genome-wide pQTL analysis of protein expression regulatory networks in the human liver. BMC Biol. 2020 Aug 10;18(1):97.

84. Robins C, Liu Y, Fan W, Duong DM, Meigs J, Harerimana NV, et al. Genetic control of the human brain proteome. Am J Hum Genet. 2021 Mar 4;108(3):400–10.

